# SynaptoTagMe: A Toolkit for *In Vivo* Mapping and Modulating Neurotransmission at Single-Cell Resolution

**DOI:** 10.1101/2025.08.18.670838

**Authors:** Andrea Cuentas-Condori, Patricia Chanabá-López, Matthew Thomas, Likui Feng, Aaron Wolfe, Peter Agoba, Matthew L. Schwartz, Maximillian Brown, Margaret Ebert, Erik Jorgensen, Cornelia I. Bargmann, Daniel Colón-Ramos

## Abstract

Understanding the organization and regulation of neurotransmission at the level of individual neurons and synapses requires tools that can track and manipulate transmitter-specific vesicles *in vivo*. Here, we present SynaptoTagMe, a suite of genetic tools in *Caenorhabditis elegans* to fluorescently label and conditionally ablate the vesicular transporters for glutamate, GABA, acetylcholine, and monoamines. Using a structure-guided approach informed by protein topology and evolutionary conservation, we engineered endogenously tagged versions for each transporter that maintain their physiological function while allowing for cell-specific, bright, and stable visualization. We also developed conditional knockout strains that enable targeted disruption of neurotransmitter synthesis or packaging in single neurons. We applied this toolkit to map co-expression of vesicular transporters across the *C. elegans* nervous system, revealing that over 10% of neurons exhibit co-transmission. Using the ADF sensory neuron as a case study, we demonstrate that serotonin and acetylcholine are trafficked in partially distinct vesicle pools. Our approach provides a powerful platform for mapping, monitoring, and manipulating neurotransmitter identity and use *in vivo*. The molecular strategies described here are likely applicable across species, offering a generalizable approach to dissect synaptic communication *in vivo*.

## INTRODUCTION

Understanding how the nervous system generates behavior requires tools that can resolve the molecular identity, spatial localization, and functional contribution of neurotransmitters *in vivo*. Neurotransmitters are the primary means by which neurons communicate, and their synthesis, packaging, and release are governed by evolutionarily conserved molecular pathways shared from *Caenorhabditis elegans* to vertebrates (Südhof, 2021). These transmitters shape the strength, kinetics (tonic vs. phasic), and polarity (excitatory vs inhibitory) of synaptic transmission, thereby influencing how information is processed and how behavior is regulated (Crawford & Kavalali, 2015; Gouwens et al., 2020; Kamalova & Nakagawa, 2021; Liu et al., 2021). Because neurotransmitters are central to defining the functional properties of synapses, understanding their identity and dynamics is essential for interpreting circuit function. Even in organisms with complete connectomes—such as *C. elegans* (White et al., 1986) and *Drosophila melanogaster* (Scheffer et al., 2020; Seggewisse & Winding, 2024; Yi et al., 2024) anatomical connectivity alone cannot explain how neural circuits generate behaviors. To build accurate, testable models of circuit function, it is necessary to also determine which neurotransmitters are used at specific synapses and how their release is spatiotemporally organized and regulated *in vivo*. Yet, despite the centrality of neurotransmitters to circuit logic, the field still lacks broadly applicable tools to visualize and manipulate transmitter-specific vesicle pools with the precision needed to study their roles in intact, living animals.

Traditional approaches—such as in situ hybridization, immunohistochemistry, and transcriptomics—have been instrumental in mapping neurotransmitter identity. These methods, however, often lack cell-specific control, temporal resolution, or the ability to monitor transmitter usage dynamically within intact circuits. Moreover, neurotransmitter identity can change in response to environmental or physiological cues. For example, neurons may co-release multiple transmitters or modulate transmitter usage depending on stress, activity, or developmental stage, and these changes have consequences in animal behavior and circuit function (Chen et al., 2023; Li et al., 2024; Maddaloni et al., 2024; Sitko et al., 2025; Wu et al., 2020). Tracking and manipulating these physiological changes require new tools that allow endogenous, live imaging and functional interrogation of neurotransmitters in single neurons.

Vesicular transporters offer a strategic entry point for such investigations. These multi-pass membrane proteins package specific neurotransmitters—such as glutamate, GABA, acetylcholine, and monoamines—into synaptic vesicles and are necessary and sometimes sufficient for defining a neuron’s transmitter phenotype (Edwards, 2007). Because they are genetically encoded and highly conserved (Alfonso et al., 1993; Bellocchio et al., 1998; Chaudhry et al., 1998; Lee et al., 1999; McIntire, Jorgensen, & Horvitz, 1993; McIntire et al., 1997; Roghani et al., 1994), vesicular transporters provide a powerful molecular handle for developing generalizable tools that probe synaptic identity and function across species. Tagging these transporters can offer direct, real-time readouts of presynaptic signaling and enable manipulations that dissect the functional contribution of specific neurotransmitters *in vivo* (Li et al., 2020). Yet for these tools to be broadly useful, it is essential that tagging does not disrupt the localization or function of the transporter. If appropriate insertion sites can be identified and validated functionally, the evolutionary conservation of vesicular transporters suggests that such designs could serve as generalizable platforms across systems and species.

Here, we present SynaptoTagMe, a comprehensive toolkit for tracking and manipulating transmitter-specific vesicles in *C. elegans*. Using a structure-guided approach informed by predicted protein topology and sequence conservation, we engineered endogenously tagged versions of the vesicular transporters for glutamate (EAT-4/VGLUT), GABA (UNC-47/VGAT), acetylcholine (UNC-17/VAChT), and monoamines (CAT-1/VMAT). We validated *in vivo* that the tagged transporters retain functionality and enables bright, cell-specific imaging. In parallel, we developed conditional knockout strains that enabled spatiotemporal access to the ablation of the packaging or synthesis of specific neurotransmitters in defined neurons, allowing causal tests of neurotransmitter function at the single-cell level within behaving animals.

We applied this toolkit to identify neurons that co-express multiple vesicular transporters, revealing that 10% of *C. elegans* neurons contain the machinery for co-transmission. Focusing then on the ADF sensory neuron, we validate that ADF expresses the machinery for co-transmission of serotonin and acetylcholine. We demonstrate that serotonin and acetylcholine are packaged in partially distinct vesicle populations. Together, our observations suggest that co-transmission can be spatially organized, offering a refined view of how individual neurons diversify their signaling output *in vivo*. Our discoveries also highlight co-transmission as a widespread and previously underappreciated feature of nervous system organization, rather than a rare or specialized exception. Co-transmission is not unique to *C. elegans*; in *Drosophila* the VAChT protein can be modulated in GABAergic and glutamatergic neurons by microRNAs (Chen et al., 2023); in mammals, serotonergic neurons in the dorsal raphe co-release glutamate or GABA depending on context (Li et al., 2024), while starburst amacrine cells in the retina release both acetylcholine and GABA with distinct calcium sensitivities (Lee et al., 2010; Morrie & Feller, 2015). These examples, along with our findings, underscore the evolutionary conservation of co-transmission as a mechanism for expanding the functional repertoire of single neurons.

By enabling simultaneous visualization of different transmitter-specific vesicle pools within the same neuron, our tools uncover molecular heterogeneity at individual synapses and reveal new layers of synaptic plasticity. More broadly, our findings establish a functional framework for probing neurotransmitter dynamics, synaptic architecture, and co-transmission *in vivo*. The strategies developed here are generalizable to other model systems and open new avenues for dissecting neural circuit logic with molecular and cellular precision.

## RESULTS

### A systematic strategy for tagging and manipulating transmitter-specific vesicles *in vivo*

All synaptic vesicle transporters are multi-pass transmembrane proteins with structural loops facing either the cytosolic or luminal space. To visualize transmitter-specific vesicle pools *in vivo*, we developed a suite of fluorescently tagged, functional versions of the vesicular transporters for glutamate (EAT-4/VGLUT), GABA (UNC-47/VGAT), acetylcholine (UNC-17/VAChT), and monoamines (CAT-1/VMAT) in *C. elegans*. We chose these four neurotransmitter classes because they are used by more than 90% of the neurons in *C. elegans* (Wang et al., 2024). We used a systematic design pipeline that integrated (1) protein topology predictions, (2) evolutionary conservation, and (3) structure-guided fluorophore placement to identify regions of each transporter suitable for tagging without disrupting function. These approaches were used to generate endogenous knock-in alleles with bright, cell-specific labeling through Flippase recombinase systems (Schwartz & Jorgensen, 2016) or self-assembling split-GFP tags (He et al., 2019) (Figure 1). When possible, tools were developed for both green and red-based fluorophores to allow for multi-color imaging. For each transporter, we also created matched conditional knockout strains by inserting FRT-flanked cassettes to disrupt neurotransmitter packaging or synthesis in defined cells, adding to the existing cell-specific knockout tools available in the field (Huang et al., 2023; Lopez-Cruz et al., 2019) (Figure 1). To drive expression of Flippase in GABAergic and Cholinergic neurons we inserted Flippase into the *unc-47* and *unc-17* locus after a self-cleaving 2A peptide sequence (Ahier & Jarriault, 2014), and used available Flippase drivers for Serotonergic and Dopaminergic neurons (Muñoz-Jiménez et al., 2017). Together, these new tools allow precise labeling and *loss-of-function* analysis of transmitter-specific vesicles in intact circuits and behaving animals (summarized in Table 1). All strains created in this study are accessible upon request from the Caenorhabditis Genetics Center (CGC) and the respective sequences are available at https://www.intralab.app/research-papers/cuentas-condori_etal-2026.

**Figure 1 –.**
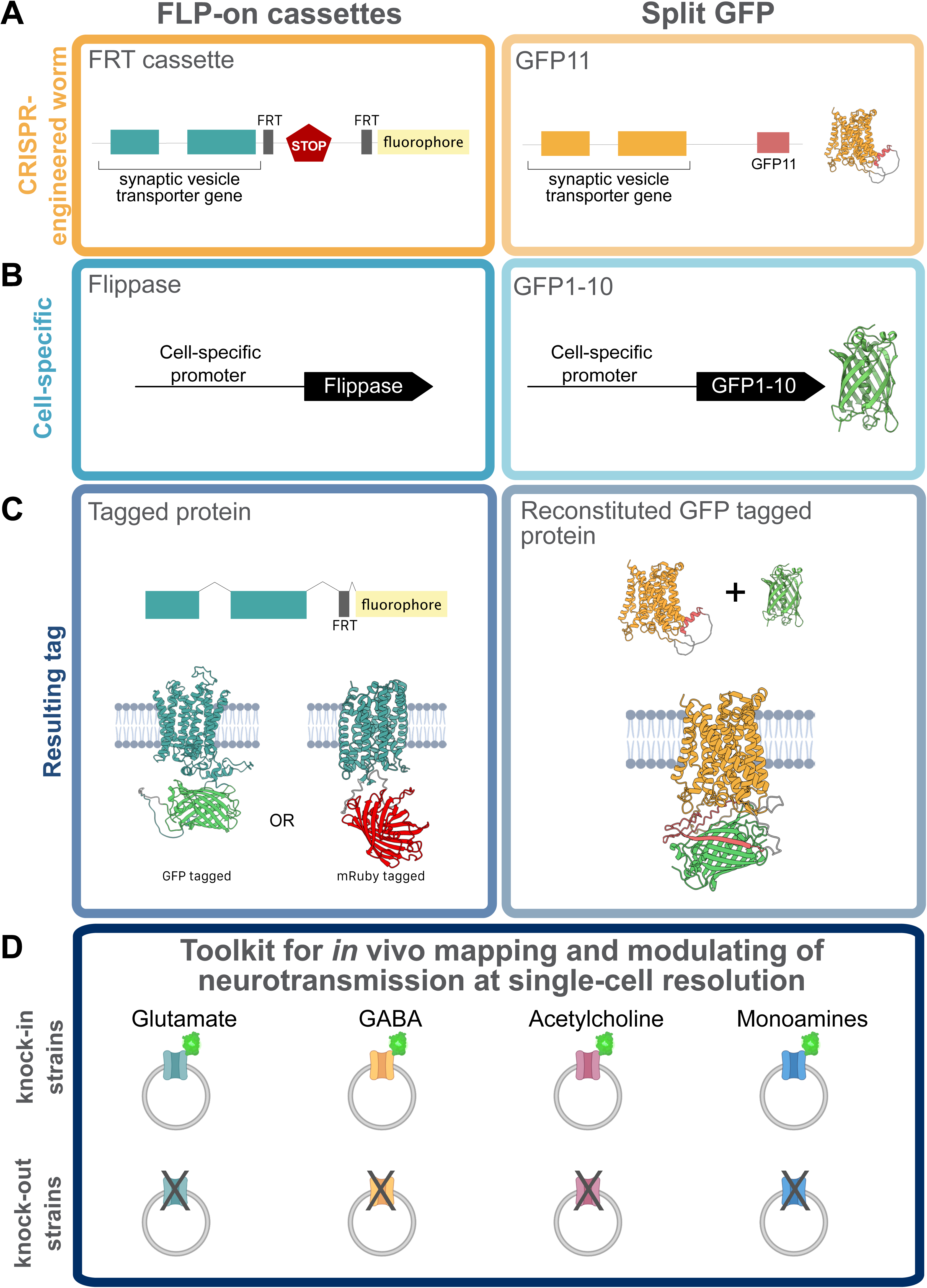
Endogenous fluorescence labeling of synaptic vesicle transporters. (Left) FLP-on and (Right) split GFP strategies to endogenously label synaptic vesicle transporters. **(A)** Cartoon representation of the CRISPR-engineered worm in the endogenous locus of the target synaptic vesicle gene. **(B)** Cell-specific driver that expresses (Left) Flippase or (Right) GFP1-10. **(C)** Resulting tagged synaptic vesicle proteins with a (Left) full-length GFP or mRuby3 or (Right) by reconstituting GFP in the cell of interest. **(D)** Schematic of the resulting toolkit to label and eliminate the endogenous machinery that packages or synthesizes glutamate, GABA, acetylcholine and monoamines.

**Table 1.**
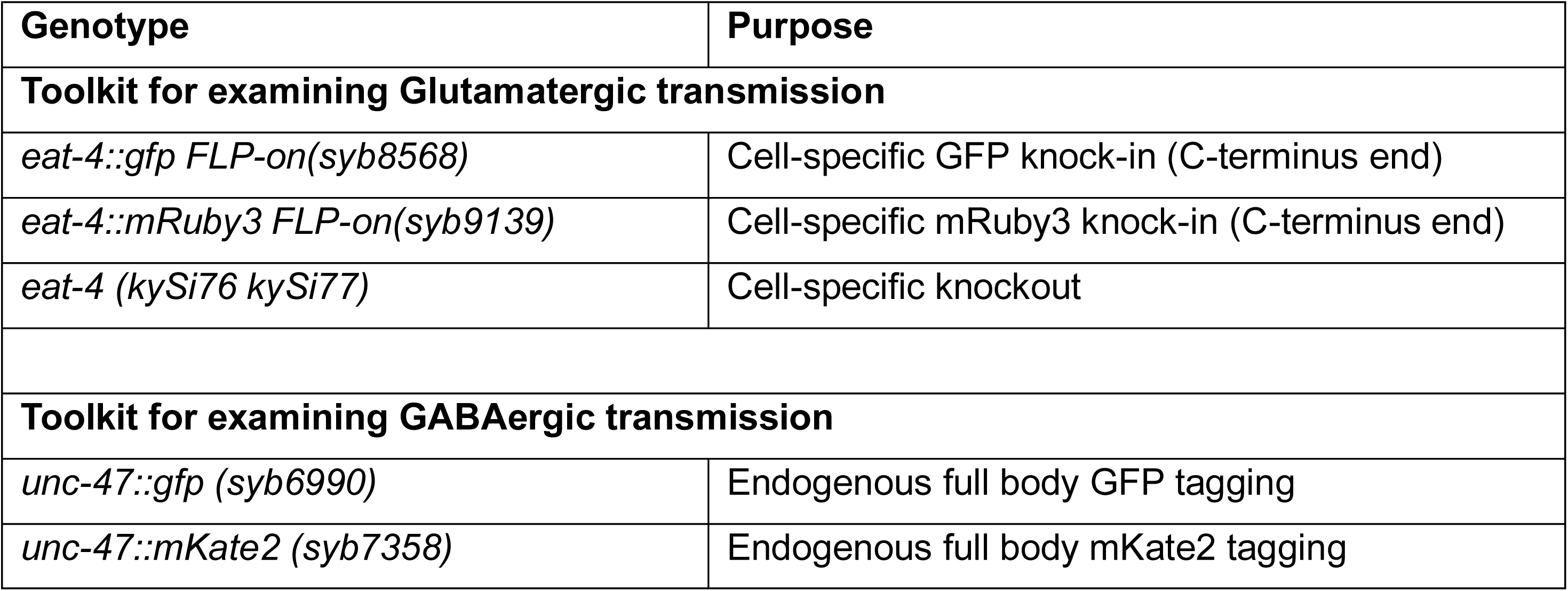
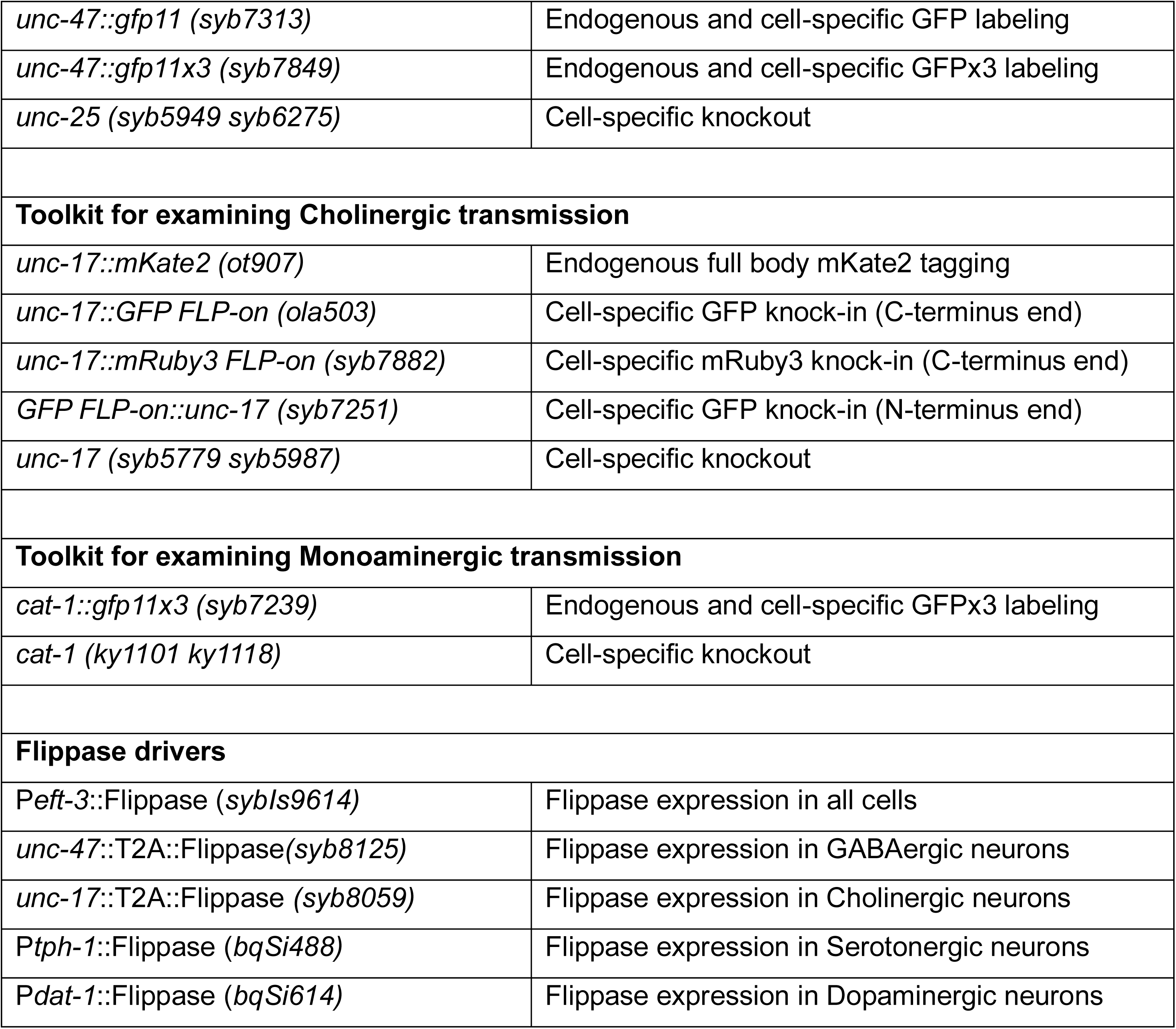
Cellular, genetic and molecular tools to probe neurotransmission in single cells.

| Genotype | Purpose |
| --- | --- |
| **Toolkit for examining Glutamatergic transmission** |  |
| *eat-4::gfp FLP-on(syb8568)* | Cell-specific GFP knock-in (C-terminus end) |
| *eat-4::mRuby3 FLP-on(syb9139)* | Cell-specific mRuby3 knock-in (C-terminus end) |
| *eat-4 (kySi76 kySi77)* | Cell-specific knockout |
| **Toolkit for examining GABAergic transmission** |  |
| *unc-47::gfp (syb6990)* | Endogenous full body GFP tagging |
| *unc-47::mKate2 (syb7358)* | Endogenous full body mKate2 tagging |
| *unc-47::gfp11 (syb7313)* | Endogenous and cell-specific GFP labeling |
| *unc-47::gfp11x3 (syb7849)* | Endogenous and cell-specific GFPx3 labeling |
| *unc-25 (syb5949 syb6275)* | Cell-specific knockout |
| **Toolkit for examining Cholinergic transmission** |  |
| *unc-17::mKate2 (ot907)* | Endogenous full body mKate2 tagging |
| *unc-17::GFP FLP-on (ola503)* | Cell-specific GFP knock-in (C-terminus end) |
| *unc-17::mRuby3 FLP-on (syb7882)* | Cell-specific mRuby3 knock-in (C-terminus end) |
| *GFP FLP-on::unc-17 (syb7251)* | Cell-specific GFP knock-in (N-terminus end) |
| *unc-17 (syb5779 syb5987)* | Cell-specific knockout |
| **Toolkit for examining Monoaminergic transmission** |  |
| *cat-1::gfp11x3 (syb7239)* | Endogenous and cell-specific GFPx3 labeling |
| *cat-1 (ky1101 ky1118)* | Cell-specific knockout |
| **Flippase drivers** |  |
| *Peft-3::Flippase (sybIs9614)* | Flippase expression in all cells |
| *unc-47::T2A::Flippase(syb8125)* | Flippase expression in GABAergic neurons |
| *unc-17::T2A::Flippase (syb8059)* | Flippase expression in Cholinergic neurons |
| *Ptph-1::Flippase (bqSi488)* | Flippase expression in Serotonergic neurons |
| *Pdat-1::Flippase (bqSi614)* | Flippase expression in Dopaminergic neurons |

### Functional labeling of glutamatergic vesicles via EAT-4/VGLUT

Glutamate functions as a key excitatory neurotransmitter in the nervous system, and its packaging into synaptic vesicles requires the conserved Vesicular Glutamate Transporter (VGLUT) (Bellocchio et al., 1998; Lee et al., 1999), which is sufficient to confer glutamatergic identity to a neuron. In *C. elegans*, the VGLUT homolog EAT-4 is expressed in 43 of the 118 neuronal classes catalogued (Wang et al., 2024). EAT-4/VGLUT is predicted to have 12 transmembrane domains (Figure 2A), and prior tools have allowed for cell-specific knockout of its full coding sequence (Lopez-Cruz et al., 2019). A previously reported transgene with EAT-4/VGLUT fused to GFP demonstrated localization to synapses (Yu & Chang, 2022); here we extend this approach to an endogenously tagged allele.

**Figure 2 –.**
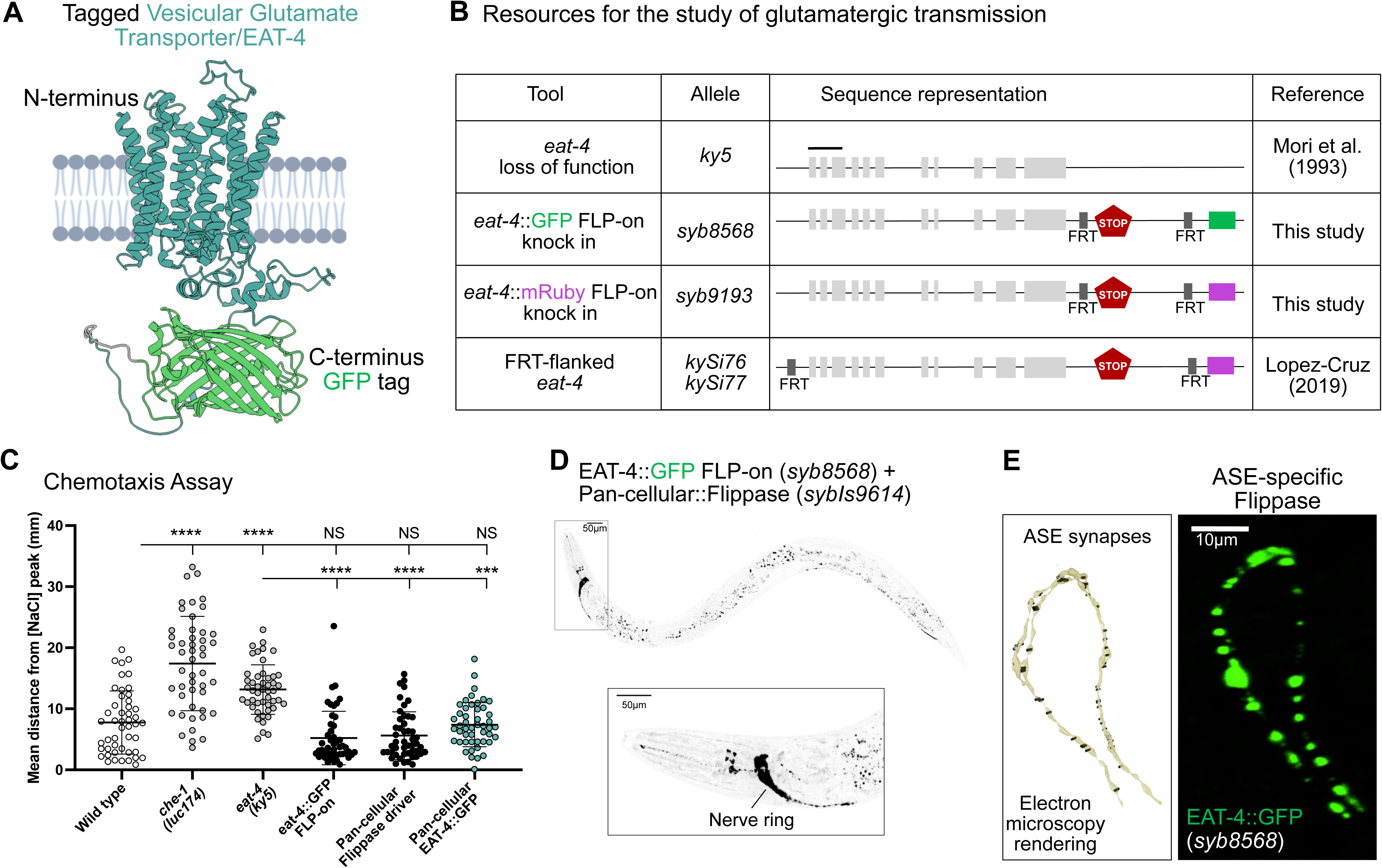
Probing glutamatergic transmission in *C. elegans*. **(A)** Predicted protein structure (using AlphaFold Protein Structure Database) of EAT-4 (cyan) tagged with GFP (green) (Abramson et al., 2024). The last amino acid in the C-terminus end (W576) corresponds to the tagged residue. **(B)** Schematic of the Vesicular Glutamate Transporter (VGLUT/*eat-4*) gene structure, loss of function allele (*ky5*) and endogenously tagged versions built for this study and by others. *eat-4* (*ky5*) mutants lack the first three exons. For cell-specific knock-in (KI) tools, GFP (*syb8568*) and mRuby (*syb9193*) FLP-ON cassettes (Schwartz & Jorgensen, 2016) were inserted before the STOP codon. To cell-specifically knock out (KO) *eat-4*, two FRT sites flank the coding sequence of the *eat-4* gene (*kySi76 kySi77*) (Lopez-Cruz et al., 2019). When recombination takes place, the *eat-4* coding sequence is removed and cytosolic mCherry is inserted in-frame to be expressed as a proxy for *eat-4* sequence removal. **(C)** Mutations in the *che-1* (prevents ASE development (Uchida et al., 2003)) (17.42 ± 7.7 mm) or *eat-4* gene (13.15 ± 4 mm) results in disrupted migration across the salt gradient. Wild-type animals (7.76 ± 5.2 mm) migrate across the salt gradient similarly to EAT-4::GFP FLP-on (*syb*8568) animals that express (7.6 ± 3.6 mm) or not (4.9 ± 4.3 mm) flippase pan-cellularly (*sybIs9614*). Animals that express flippase in all cells (5.61 ± 3.9 mm) migrate across the salt gradient as a wild-type animals. Results represent the mean distance of each worm from the salt peak, averaged across the final minute of the assay, with each dot representing a single animal. Plots are overlaid with Mean ± Standard Deviation. Kruskal-Wallis test with Dunn’s multiple comparison post hoc test. **** represents p<0.0001; *** represents p<0.001 and NS means “not significant”. **(D)** (Top) Fluorescent image of an adult worm expressing endogenously labeled EAT-4 with GFP in all cells (P*eft-3*::Flippase). Scale Bar = 50μm. (Bottom) Zoom-in area of the head shows EAT-4 expressed predominantly in the nerve ring. Scale Bar = 10μm. **(E)** (Left) Electron microscopy rendering of ASE synapses in an L4 wild-type animal (White et al., 1986) (image generated with NeuroSC (Koonce et al., 2025). (Right) Fluorescent image of endogenously tagged EAT-4 protein cell-specifically in the ASE axons (P*flp-6*::Flippase). Scale Bar = 10μm.

To generate a bright, functional reporter that reflects endogenous EAT-4/VGLUT localization *in vivo*, we inserted a GFP tag into the protein C-terminal cytoplasmic domain using a FLP-on cassette (Schwartz & Jorgensen, 2016). The C-terminus was chosen based on conservation analysis and AlphaFold structural predictions (Jumper et al., 2021; Pei & Grishin, 2001), which identified it as a cytosolic and weakly conserved region, minimizing the risk of disrupting conserved protein functions (Figure S1A-B). GFP was inserted just before the STOP codon (Figure 2B). To examine if introduction of GFP into the endogenous EAT-4/VGLUT gene affected function, we assessed NaCl chemotaxis (Figure S1C) —an EAT-4–dependent learning behavior mediated by the ASE neurons (Figure S1C-D) (Bargmann & Horvitz, 1991; Sato et al., 2021; Uchida et al., 2003). EAT-4::GFP FLP-on strains, with or without Flippase expression in all neurons or selectively in ASE neurons, displayed normal chemotaxis behavior (Figure 2C and S1D), in contrast to *eat-4(ky5)* loss-of-function mutants or mutant animals with defects in chemosensory neurons, including ASE (*che-1* mutants; Figure 2C, S1C-D). Our findings suggest that the tagged protein remains functional and capable of sustaining known glutamate-dependent behaviors in the organism.

The insertion of the FLP-on cassette enables expression of EAT-4/VGLUT::GFP upon cell-specific expression on the FLP recombinase. To validate its use, we expressed pan-cellular flippase via the *eft-3* promoter (Seydoux & Fire, 1994) in the worms engineered with the EAT-4::GFP FLP-on cassette. We observed bright EAT-4::GFP signal throughout the nervous system, especially in the nerve ring and sensory neurons (Figure 2D), consistent with earlier transcriptional reporters of the *eat-4* gene (Lee et al., 1999; Serrano-Saiz et al., 2013). Moreover, when Flippase was driven specifically in ASE neurons (using the ASE-specific promoter, P*flp-6*), we observed punctate labeling along ASE axons, matching the distribution of presynaptic sites identified by serial electron microscopy (Figure 2E), and cataloged in NeuroSC (Koonce et al., 2024; White et al., 1986). Consistent with this, we observed that endogenous GFP::RAB-3 in ASE similarly localizes in a punctate pattern along the axons in a pattern that is reminiscent to that seen for endogenous EAT-4::GFP (Figure S2 and Figure 2E).

To expand the utility of this tool for multicolor imaging, we also generated a red-shifted FLP-on reporter for the *eat-4/VGLUT* gene by inserting mRuby3 (Bajar et al., 2016) with a *C. elegans*-optimized sequence at the same C-terminal site (Figure 1B). These spectrally distinct reporters, when combined with the previously developed *eat-4* conditional knockout (Lopez-Cruz et al., 2019) provide a comprehensive toolkit for dissecting glutamatergic transmission in a cell-specific manner *in vivo*.

### Cell-specific imaging and silencing of GABAergic neurotransmission

GABAergic neurons package GABA into synaptic vesicles via the conserved vesicular GABA transporter VGAT(Chaudhry et al., 1998; McIntire et al., 1997). In *C. elegans*, the VGAT homolog UNC-47 is expressed in 11 of the 118 neuronal classes (Gendrel et al., 2016; Wang et al., 2024). Based on *in vivo* data, the N-terminus of VGAT is cytoplasmic while the C-terminus is luminal (Martens et al., 2008). The N-terminus contains dileucine motifs critical for proper trafficking (Santos et al., 2013), and to preserve transporter function we focused on tagging long cytoplasmic loops. Structural predictions from AlphaFold indicate that UNC-47 has 11 transmembrane domains (Figure 3A), and we identified the cytosolic loop between transmembrane domains 2 and 3— a long (13 amino acids) region (Figure S3A-B)—as an optimal tagging site.

**Figure 3 –.**
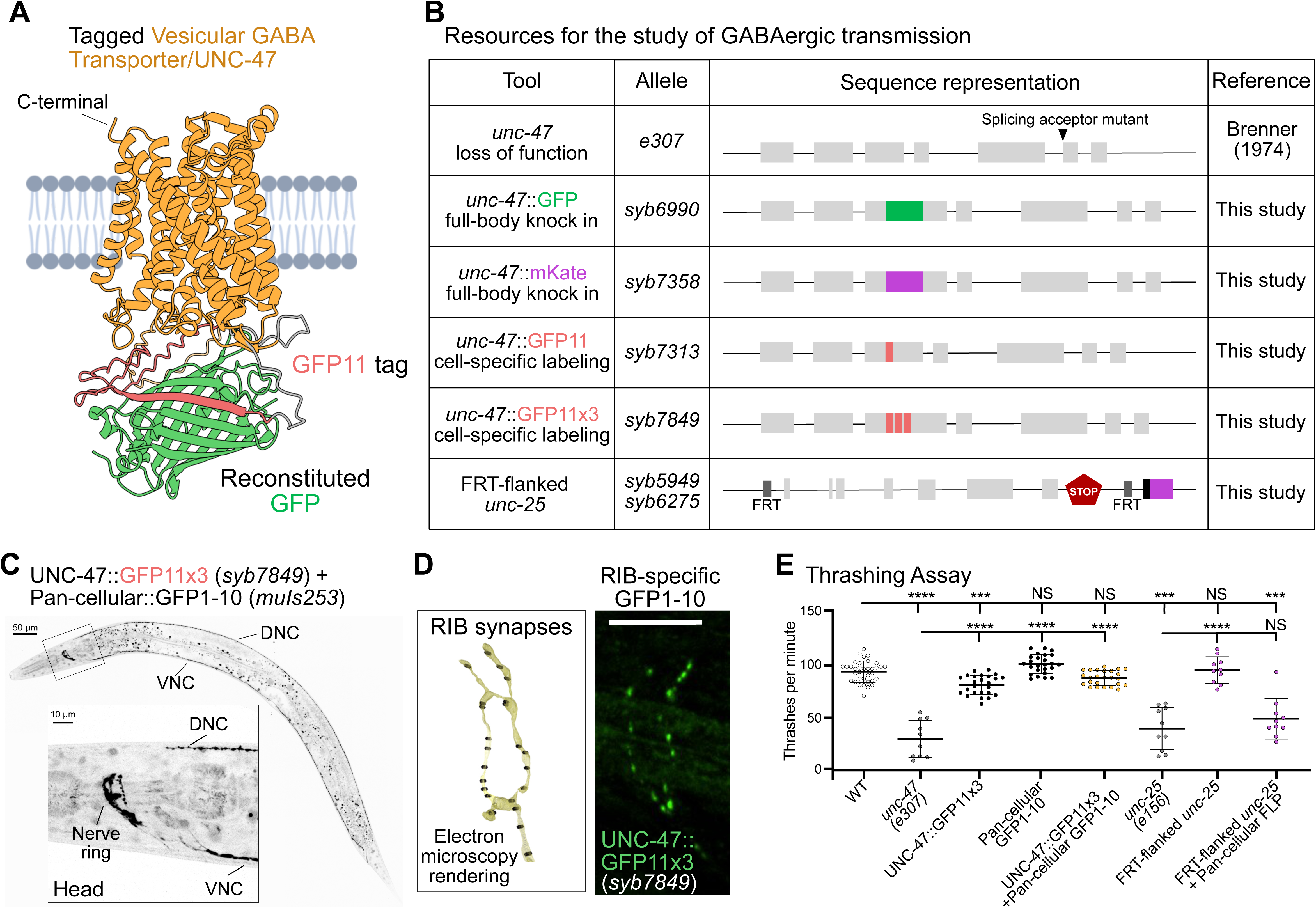
Probing GABAergic transmission in *C. elegans*. **(A)** UNC-47 predicted protein structure from AlphaFold Protein Structure Database. GFP11 tag (red) was added between amino acids E145 and N146. Complementary GFP1-10 (green) was modeled as bound to GFP11. **(B)** Schematics of the Vesicular GABA Transporter (VGAT/*unc-47*) loss of function allele and endogenously tagged versions built for this study. *unc-47(e307)* mutant animals have a single base pair substitution (G to A in the first nucleotide of exon 6) that results in a splicing acceptor mutant. Fluorescent tags GFP (*syb6990*) and mKate2 (*syb7358*) were inserted between amino acids E145 and N146. For cell-specific endogenous labeling, UNC-47 was tagged with one (*syb7313*) or three copies (*syb7849*) of GFP11. To silence GABA transmission, we flanked the Glutamic Acid Decarboxylase/*unc-25* coding sequence with two FRT sites (*syb5949 syb6275*). Upon recombination, the *unc-25* coding sequence is removed and nuclear (black) mCherry (purple) is designed to be in-frame and expressed as a proxy for *unc-25* sequence removal. **(C)** (Top) Fluorescent image of an adult worm expressing endogenously labeled UNC-47 with reconstituted split-GFP in all cells (P*eft-3*::GFP1-10). Scale Bar = 50μm. (Bottom) Zoom-in area of the head shows UNC-47 expressed in the nerve ring and nerve cords. Scale Bar = 10μm. **(D)** (Left) Electron microscopy renderings of RIB synapses in an L4 wild-type animal (White et al., 1986) (image generated with NeuroSC (Koonce et al., 2025)). (Right) RIB-specific UNC-47 puncta (green) *in-vivo* using the UNC-47:GFP11 with reconstituted split-GFP in RIB (P*sto-3b*::GFP1-10) Scale Bar = 10μm. **(E)** *unc-47(e307)* mutant animals thrash significantly less per minute (30.8 ± 17) than wild-type animals (94.5 ± 10). Animals with non-reconstituted UNC-47::GFP11X3 (82.2 ± 9) (*syb7849*) thrash slightly less than wild-type animals (94.5 ± 10), while animals that express pan-cellular GFP1-10 (100.5 ± 9) (*muIs253*) are no different than wild-type animals. UNC-47::GFP11x3 animals that express Pan-cellular::GFP1-10 (with reconstituted GFP) (88.6 ± 7) thrash similarly to wild-type animals. FRT-flanked *unc-25* animals (*syb5949 syb6275*) that do not express Flippase (95.8 ± 12) thrash similarly to wild-type animals (91 ± 8). Cell-specific *unc-25* knockout animals (*syb5949 syb6275*) that express Flippase in every neuron (*bqSi506*) (49.36 ± 19) thrash to the same extent as *unc-25* (*e156*) mutant animals (40.2 ± 19). Plots are overlaid with Mean ± Standard Deviation. Kruskal-Wallis test with Dunn’s multiple comparison post hoc test. **** represents p<0.0001; *** represents p<0.001; and NS means “not significant”.

Within this loop, AlphaFold predicts two beta-sheet regions with high confidence. We inserted GFP between amino acids E145 and N146, immediately following the first predicted beta sheet, to avoid disrupting secondary structures (Figure 3A, S3A-B). To assess functionality of the newly engineered UNC-47/VGAT::GFP strain, we performed thrashing assays on *unc-47 (e307)* mutants, which show impaired locomotion due to loss of GABA signaling at neuromuscular junctions (McIntire, Jorgensen, Kaplan, et al., 1993). Expression of UNC-47::GFP from an extrachromosomal array rescued the thrashing defect to wild-type levels (Figure S3C and D), as expected. We next generated endogenous knock-ins of GFP and mKate2 at the same site but in the endogenous *unc-47* locus, and observed wild-type locomotion for these strains, consistent with the insertion of the fluorophores not affecting endogenous function of the transporter (Figure S3D). These strains showed bright, punctate fluorescence in the nerve ring and along the dorsal and ventral nerve cords (Figure S3E-F), consistent with previously reported *unc-47* expression patterns (McIntire et al., 1997). Together, these results demonstrate that inserting a fluorescent protein at position E145 results in a functionally tagged UNC-47/VGAT reporter that enables endogenous visualization of the protein.

To enable *in vivo* visualization of GABAergic vesicles in single-cells, we next generated two UNC-47::splitGFP alleles by inserting either one or three tandem copies of GFP11 at the E145 position (Figure 3B). We used the splitGFP approach to avoid disruptions of the protein structures due to the introduction of the FRT-cassettes at an internal sequence site. By leveraging the self-assembling property of the GFP beta barrel, knock-in of the eleventh beta strand (GFP11) results in labeling of a protein that is only visible when the complementary GFP1-10 is expressed in the same cell. This property results in a combinatorial labeling strategy, in which cell-specific labeling is achieved only in those cells that express both the GFP11 and the GFP 1-10 (He et al., 2019). To validate these tools, we first achieved pan-cellular expression of GFP1-10 (*eft-3* promoter) (Seydoux & Fire, 1994) in animals carrying the UNC-47::GFP11x3 *(syb7849)* allele. We observed GABAergic synapses throughout the nerve ring and nerve cords (Figure 3C), similar to full-body knock-in strains (Figure S3E-F). To then visualize GABAergic vesicles in subsets of cells, we expressed GFP1-10 in the GABAergic DD motor neurons using the *flp-13* promoter and in animals carrying the GFP11 (*syb7313*) or UNC-47::GFP11x3 *(syb7849)* alleles. We observed punctate reconstituted signal in the dorsal nerve cord of both GFP11 and GFP11x3 strains, consistent with the known distribution of DD synapses (Figure S4E). The triple GFP11 version produced significantly brighter signal (Figure S4E-G), in line with reports of enhanced fluorescence from multimerized tags (He et al., 2019).

To then visualize GABAergic synapses in single cells, we expressed GFP1-10 under an RIB-interneuron, cell-specific promoter. We selected RIB because it is a GABAergic interneuron embedded in the nerve ring and proximal to three other GABAergic interneurons with overlapping neurites that impede visualization of RIB-specific synapses *in vivo* when using traditional approaches (Figure S4A-C). Reconstituted fluorescence using our tools enabled visualization of RIB-specific synapses, and the observed synaptic pattern was consistent with the pattern expected from electron microscopy reconstructions (Figure 3D) (Koonce et al., 2024; White et al., 1986) and from expression of endogenous mScarlet::RAB-3 in RIB neurons (Figure S4D). These findings underscore the value of the tool in labeling GABAergic synapses in individual cells *in vivo*.

To functionally dissect GABAergic transmission, we developed a conditional knockout of *unc-25*, the gene encoding Glutamic Acid Decarboxylase (GAD), which catalyzes GABA synthesis. We decided to target *unc-25*/GAD because it results in the elimination of GABA from neurons (Gendrel et al., 2016). We flanked the *unc-25* coding sequence with FRT sites and inserted a nuclear mCherry reporter downstream to indicate successful recombination (Figure 3B). We then validated the tool by using thrashing assays. Consistent with previous findings, *unc-25* null mutants show severely reduced locomotion in the thrashing assays (McIntire, Jorgensen, Kaplan, et al., 1993) (Figure 3E). We observed that animals carrying the conditional knockout allele behaved normally in the absence of Flippase, but that pan-cellular Flippase expression in all GABAergic neurons (*unc-47* promoter), which is expected to result in loss of GABAergic neurotransmission, phenocopied the thrashing defect of *unc-25* mutants (Figure 3E). These results confirm that this tool effectively eliminates *unc-25/*GAD activity, with the capacity to be activated in a cell-specific manner and allows investigation of how GABAergic transmission contributes to neural function and behavior.

### Functional labeling of cholinergic vesicles via UNC-17/VAChT

The cholinergic identity of neurons is defined by the expression of a conserved gene locus, shared from nematodes to vertebrates, that includes both the acetylcholine synthesis enzyme Choline Acetyltransferase (ChAT) and the Vesicular Acetylcholine Transporter (VAChT) (Eiden, 1998). In *C. elegans*, the VAChT homolog UNC-17 is expressed in 57 of the 118 neuronal classes (Wang et al., 2024). Based on prior studies, both the N- and C-termini of VAChT face the cytosol and contain regulatory motifs important for trafficking (Fei et al., 2008). AlphaFold predicts UNC-17 has 12 transmembrane domains, but most cytosolic loops are very short (<5 amino acids), except for the third cytoplasmic loop, which contains 28 amino acids and exhibits relatively high sequence conservation (Figure S5A-B).

To determine suitable tagging sites for imaging UNC-17 without disrupting function, we tested three locations: the conserved third cytosolic loop (site 1), the N-terminus (site 2), and the C-terminus (site 3) (Figure S5B-C and S5E). We used a thrashing assay to assess function, as *unc-17(e245)* null mutants fail to thrash in liquid and are rescued by the re-expression of untagged UNC-17 under its own promoter (Figure S5F). We generated transgenic strains with a rescue array containing the *unc-17* gene with GFP inserted into these three sites. We observed that GFP insertion into site 1 failed to rescue the phenotype, suggesting disruption of UNC-17 function. In contrast, tagging either the N-terminus or C-terminus restored normal thrashing (Figure S5E–F), indicating that these positions tolerate modification. Informed by these rescue experiments, we then generated two FLP-on conditional knockout alleles (Schwartz & Jorgensen, 2016) with GFP inserted at either the N-terminus (*syb7251*) or C-terminus (*ola503*) (Figure 4A-B). To test whether the FLP-on cassettes affected protein function, we examined behavior before Flippase expression and found that both alleles behaved like wild-type animals (Figure S5F). Because splicing regulatory elements are located near the 5’ end of the *unc-17* gene and are required for successful splicing of the cholinergic locus (both *unc-17* and *cha-1* transcripts) (Mathews et al., 2015), we proceeded with the C-terminally tagged *ola503* allele, which leaves the 5’ region intact. We tested whether pan-cellular expression of Flippase in this strain impairs behavior. Animals with global GFP-tagged UNC-17 showed wild-type thrashing in liquid (Figure 4C), confirming that the tagged transporter is functional and validating this approach for cell-specific labeling of cholinergic vesicle pools.

**Figure 4 –.**
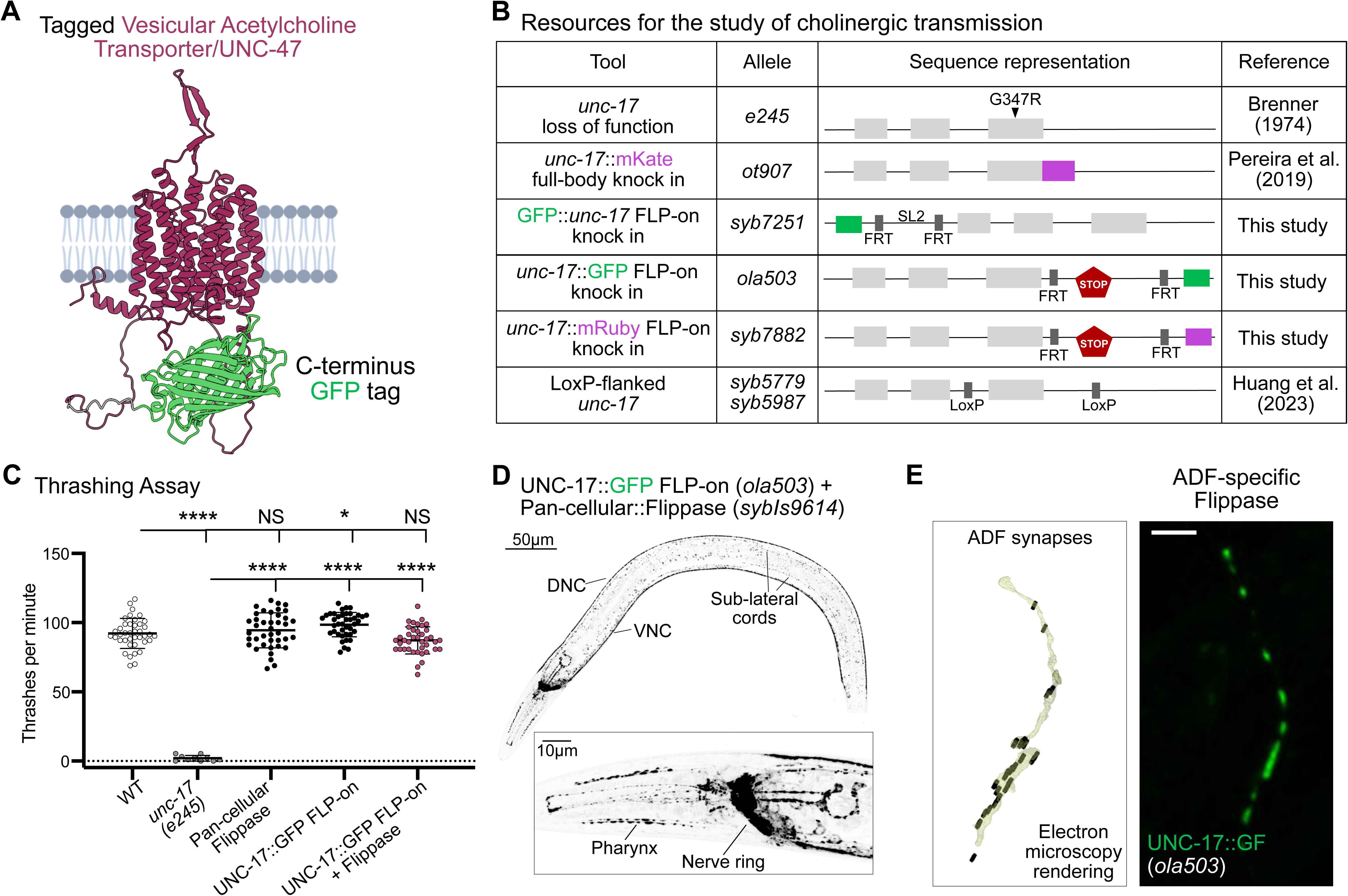
Probing cholinergic transmission in *C. elegans*. **(A)** Predicted UNC-17 protein structure (magenta) tagged with GFP (green) on the last amino acid in the C-terminus end (W532). **(B)** Schematics of the Vesicular Acetylcholine Transporter (VACHT/*unc-17*) loss-of-function allele and endogenously tagged versions tested in this study (and built by others). *unc-17(e245)* mutant animals have a single base pair substitution in the third exon which leads to an amino acid change (arrowhead). Full-body knock-in animal labels UNC-17 with mKate (*ot907*). For cell-specific labeling, the N-terminus GFP FLP-ON cassette (*syb7251*) was inserted between amino acids P6 and V7 (See Methods). Similarly, the C-terminus GFP FLP-ON cassette (*ola503*) and mRuby FLP-on cassette (*syb7882*) were inserted before the STOP codon. **(C)** *unc-17(e245)* mutant animals (1.9 ± 2) thrash significantly less than wild-type animals (93 ± 10). UNC-17::GFP FLP-on (*ola503*) animals that express pan-cellular::Flippase (88 ± 1) (*sybIs9614*) or animals that only express pan-cellular::Flippase (94.8 ± 12) are indistinguishable from wild-type animals in their thrashing behavior. UNC-17::GFP FLP-on animals (98.7 ± 8) thrash significantly more than wild-type animals. Mean ± Standard Deviation. Brown-Forsythe ANOVA test with Dunnett’s T3 multiple comparisons post hoc test. **** represents p<0.0001; * represents p<0.05; and NS means “not significant”. **(D)** (Top) Fluorescent image of an adult worm expressing endogenously labeled UNC-17::GFP in all cells (P*eft-3*::Flippase). Scale Bar = 50μm. (Bottom) Zoom-in area of the head shows UNC-17 expressed in the nerve ring, nerve cords and sub-lateral cords. DNC = Dorsal Nerve Cord, VNC = Ventral Nerve Cord. Scale Bar = 10μm. **(E)** (Left) Electron microscopy rendering of ADF synapses in an L4 wild-type animal (White et al., 1986) (image generated with NeuroSC (Koonce et al., 2025). (Right) Fluorescence image of endogenously tagged UNC-17::GFP protein specifically in the ADF neuron. Scale Bar = 10μm.

At the cellular level, GFP labeling of UNC-17/VAChT using the *ola503* allele and pan-cellular Flippase expression (driven by the *eft-3* promoter) (Seydoux & Fire, 1994) resulted in fluorescence in the nerve ring, dorsal cord, ventral cord, and sublateral cords (Figure 4D). This expression pattern matched that of a full-body mKate2 knock-in of endogenous UNC-17/VAChT (Figure S5D) (Pereira et al., 2019) and was consistent with prior transcriptional reporters (Pereira et al., 2015) and anti-UNC-17 antibody staining (Duerr et al., 2008). The *ola503* allele also enables cell-specific labeling. When Flippase was expressed specifically in ADF neurons using the P*srh-142* promoter (Maicas et al., 2021), we observed UNC-17::GFP puncta along the ADF axon (Figure 4E). This punctate pattern aligned with known ADF presynaptic sites from electron microscopy reconstructions of L4 animals (Koonce et al., 2024; White et al., 1986). Together, these results demonstrate that the UNC-17::GFP FLP-on allele provides a reliable tool to label cholinergic vesicles in individual neurons *in vivo*, and that this labeling does not affect function in our assays. To enable multicolor imaging, we generated a red, fluorescent version of the UNC-17/VAChT tool by replacing GFP with *C. elegans*-optimized mRuby3 (Figure 4B). This mRuby-tagged allele performs as wild type animals in a thrashing assay and labels synaptic structures along the nerve ring and nerve cords (Figure S6). Combined with the previously described cell-specific *unc-17* knockout strain (Huang et al., 2023) (Figure 4B), these tools allow precise tracking and manipulation of cholinergic neurotransmission *in vivo*.

### Probing Monoaminergic Transmission

Monoamines such as serotonin, dopamine, norepinephrine, epinephrin, octopamine, and tyramine are transported into vesicles by the conserved Vesicular Monoamine transporter (Duerr et al., 1999; Erickson et al., 1992). In *C. elegans*, the VMAT homolog CAT-1 is expressed in 16 of the 118 neuronal classes (Wang et al., 2024). Recently, CAT-1 was tagged at its C-terminus with a split GFP11x3 reporter (Figure 5A–B) (Huang et al., 2023). When GFP is reconstituted pan-cellularly, this fusion produces a punctate signal enriched in the nerve ring—including the characteristic and well-known serotonergic neuron NSM—and in serotonergic neurons that are part of the reproductive organs (Figure 5C).

**Figure 5 –.**
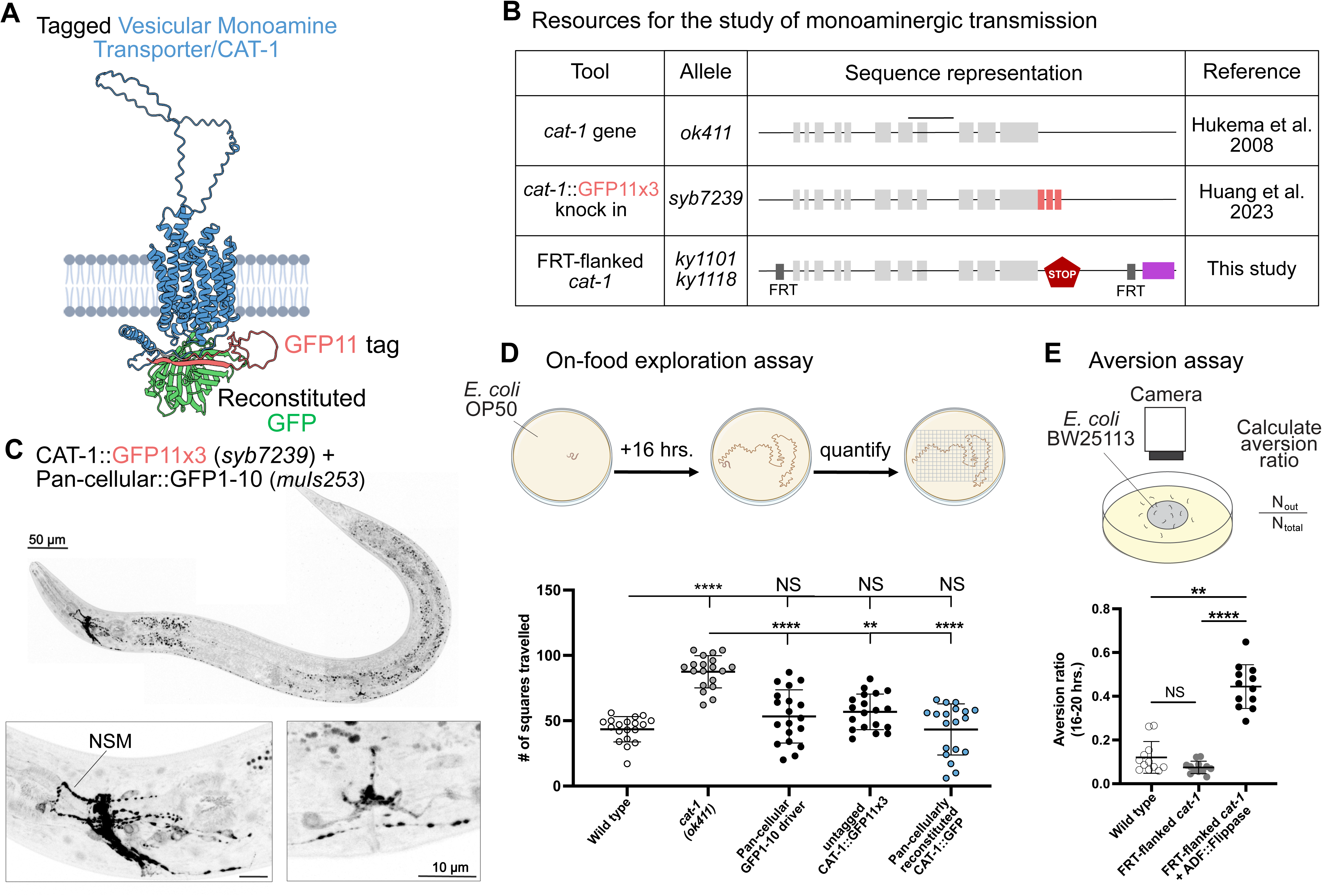
Probing Monoaminergic transmission in *C. elegans*. **(A)** Predicted CAT-1 protein structure (blue) labeled with GFP11 (red) at C-terminus end (E145). Complementary GFP1-10 (green) was modeled with bound GFP11. **(B)** Schematic of the Vesicular Monoamine Transporter (VMAT/*cat-1*) loss-of-function allele and endogenously tagged versions used to monitor monoaminergic transmission. *cat-1(ok411)* mutant animals have a deletion that spans exons 7 and 8 (black line) of the *cat-1* gene. For cell-specific labeling, three copies of GFP11 (*syb7239*) were inserted before the STOP codon. For cell-specific silencing of *cat-1* activity, two FRT sites (*ky1101 ky1118*) flank the coding sequence of the *cat-1* gene. Upon recombination, expression of cytosolic mCherry (magenta) is used as a proxy for deletion of the *cat-1* coding sequence. **(C)** (Top) Fluorescence image of reconstituted CAT-1::GFP11x3 with expression of complementary GFP1-10 in the whole animal (P*eft-3*::GFP1-10). Scale Bar = 50μm. (Bottom) Zoom-in of the (Left) head and (Right) vulva area of an adult worm shows CAT-1 expressed in the nerve ring and pharyngeal neurons as well as in the vulva. Scale Bar = 10 μm. **(D)** (Top) A well-fed day-1 adult animal is placed on NGM plates covered with a thin layer of bacteria, allowed to roam for 16 hours and the number of squares traveled was recorded. (Bottom) Wild-type animals (43.6 ± 10) roam less than *cat-1(ok411)* mutants (87.5 ± 12). CAT-1::GFP11x3 (*syb7239*) animals with GFP reconstituted (43.3 ± 20) (or not (56.8 ± 14)) roam similar to wild-type animals. Animals with pan-cellular expression of GFP1-10 (*muIs253*) (53.3 ± 20) also roam like wild-type animals. **(E)** (Top) A well-fed day-1 adult animal is placed on NGM plates covered with a thin layer of *E. coli*, allowed to roam for 20 hours and the aversion ratio was calculated. (Bottom) Wild-type animals (0.12 ± 0.07) display less aversion to *E. coli* lawns than ADF-specific *cat-1* conditional KO animals (*ky1101 ky1118; syb9159*) (0.4 ± 0.1), consistent with previous reports using serotonin-depletion mutants (Feng et al., 2025). *cat-1* conditional KO animals that do not express Flippase in ADF neurons (0.07 ± 0.03) are indistinguishable from wild-type. Mean ± Standard Deviation. Kruskal-Wallis test with Dunn’s multiple comparison post hoc test. **** represents p<0.0001; ** represents p<0.01; * represents p<0.05; and NS means “not significant”.

To determine whether the CAT-1::GFP11x3 fusion maintains protein function, we used a behavioral assay based on the role of serotonin to prevent animal exploration on a lawn of bacteria (Flavell et al., 2013). Since serotonin is packaged into vesicles by CAT-1/VMAT, similar to mutants of serotonin production (Flavell et al., 2013), *cat-1* mutants display increased exploration behavior compared to wild-type animals (Figure 5D). We found that animals with reconstituted CAT-1::GFP11x3 explore bacterial lawns at levels comparable to wild-type (Figure 5D), indicating that the GFP11x3 tag nor its reconstitution impair CAT-1 function. To complement this tool, we also developed a cell-specific *cat-1* knockout allele in which the full coding sequence is excised upon Flippase expression (Figure 5B). Removal of serotonin production (*tph-1*) specifically from ADF neurons result in increased aversion from wild type *E. coli* bacteria lawns (Feng et al., 2025). Consistent with this, cell-specific KO of *cat-1* in ADF neurons (via expression of ADF::Flippase) results in increased *E. coli* aversion (Figure 5E). Together, these tools now enable the cell-specific tracking and silencing of monoaminergic synapses in living animals.

### *In-Vivo* Identification of Co-Transmitter Neurons

Co-transmission is a conserved feature of neurons across the animal kingdom (Granger et al., 2017; Trudeau & El Mestikawy., 2018; Vaaga et al., 2014), including *C. elegans* (Duerr et al., 2008; Gendrel et al., 2016; Pereira et al., 2015; Pocock & Hobert, 2010; Serrano-Saiz et al., 2017; Serrano-Saiz et al., 2013; Taylor et al., 2021; Wang et al., 2024), but the prevalence of co-transmission *in vivo* for any given organism is not well understood. To validate the utility of our tools and to map the architecture of co-transmission in *C. elegans*, we developed an intersectional genetic strategy using Flippase recombinase and FRT-flanked fluorescent reporters (Figure 6A). We focused on identifying neurons that co-transmit glutamate or acetylcholine—the two most abundant excitatory neurotransmitters in *C. elegans*—in combination with other transmitters (Figure 6B).

**Figure 6 –.**
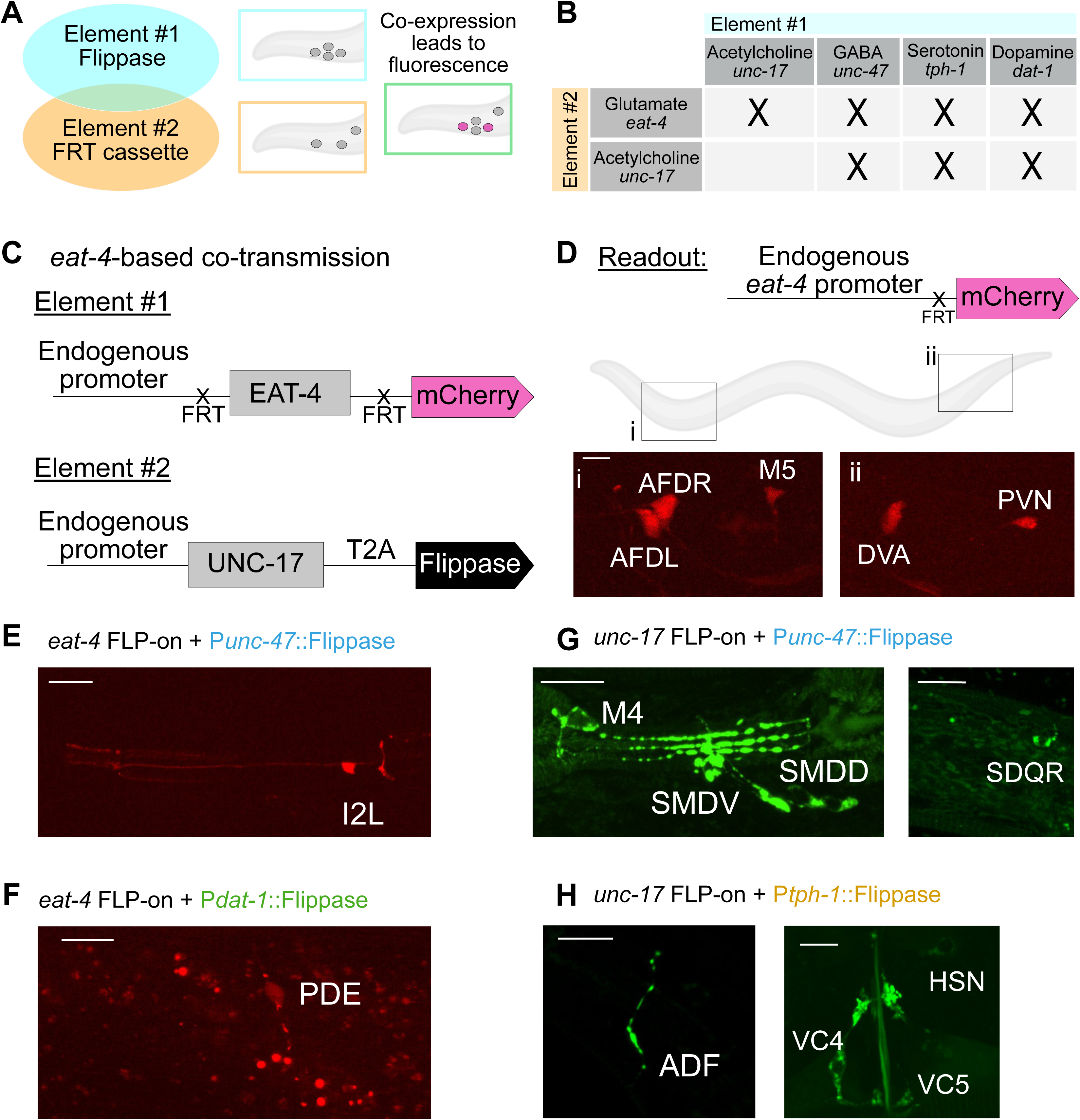
Mapping of co-transmitter neurons in the *C. elegans* nervous system. **(A)** Schematic of the approach used to find co-expression of two vesicular transporters in the same cells. Flippase drivers (Element #1, blue) and Flippase-dependent cassettes that result in fluorescence (Element #2, orange) are used. Magenta cells are seen when both elements are co-expressed (in green box). **(B)** Strategy to track the co-expression of the vesicular glutamate and acetylcholine transporter in combination with the 4 most common neurotransmitters in *C. elegans*: Acetylcholine, GABA, Serotonin and Dopamine. “Elements #1” or “Element #2” refers to the genetic elements in the schematic in Fig 6A. **(C-D)** Specific example of the genetic strategy outlined in Fig 6A, for the vesicular transporters of glutamate (EAT-4) and acetylcholine (UNC-17). **(C)** We repurposed the *eat-4* conditional KO strain (*kySi76 kySi77*) (Figure 1B) (Lopez-Cruz et al., 2019) in which the *eat-4* gene coding sequence is flanked by two FRT sites and followed by cytosolic mCherry (Element #1). We crossed this line with a strain that has an endogenously inserted self-cleaving peptide sequence (T2A) followed by Flippase before the STOP codon in the *unc-17* gene locus (Table 1). **(D)** (Top) Co-expression of EAT-4 and UNC-17 results in cytosolic mCherry. (Bottom) Cytosolic mCherry was detected in the head of three neurons: (i) AFDR, AFDL and M5; and in two neurons in the tail region: (ii) PVN, and DVA. **(E)** Fluorescence microscopy shows neurons with co-expression of **(E)** VGLUT/EAT-4 and VGAT/UNC-47; **(F)** VGLUT/EAT-4 and dopamine synthesis gene DAT-1; **(G)** VAChT/UNC-17 and VGAT/UNC-47; and **(H)** VAChT/UNC-17 and the serotonin synthesis gene TPH-1. All scale bars = 10μm.

We reasoned that if two neurotransmitters were co-expressed in the same neuron, driving Flippase under the promoter of one transmitter would activate the conditional reporter—resulting in fluorescence—only in cells also expressing a second neurotransmitter identity (Figure 6A-B). To achieve this, we used the engineered alleles for each vesicular transporter that we developed (Figures 2B, 3B, 4B, and 5B) (Table 1) and developed additional Flippase driver by modifying the locus of genes involved in the packaging of acetylcholine (*unc-17*) and GABA (*unc-47*) (Table 1). Additionally, we used available Flippase driver lines for serotonin and dopamine (Figures S7A–B, Table 1) (Muñoz-Jiménez et al., 2017).

We first used a conditional *eat-4/VGLUT* reporter strain in which cytosolic mCherry is expressed upon Flippase-mediated recombination (Lopez-Cruz et al., 2019). When Flippase was driven by the *unc-17/VAChT* promoter (cholinergic), we observed five mCherry-positive neurons in the head and tail, consistent with co-expression of *unc-17/VAChT* and *eat-4/VGLUT*. Based on cell position, neurite morphology, transcriptomic data (Taylor et al., 2021), and anatomical maps (Wang et al., 2024), we identified these neurons as AFDL, AFDR, M5, DVA, and PVN (Figure 6C–D).

Flippase expression from the GABAergic *unc-47* promoter activated *eat-4*-driven mCherry expression in a single pharyngeal neuron, identified as I2L (Figure 6E, S8A). Driving Flippase from the dopaminergic *dat-1* promoter labeled the PDE neuron (Figure 6F, S8B), while serotonergic *tph-1*-driven Flippase did not produce any mCherry-positive neurons. These results are summarized in Figure S7C.

We applied a similar strategy by using our Flippase-dependent *unc-17*::GFP reporter to identify candidate neurons that co-release acetylcholine with other neurotransmitters. In this context, when Flippase was driven from the GABAergic *unc-47* locus, we observed GFP expression in the M4, SDQR, and SMD neurons (Figure 6G, S8D). Flippase expression from the serotonergic *tph-1* promoter revealed previously described acetylcholine/serotonin co-transmitting neurons, including ADF, HSN, and VC4/VC5 (Figure 6H, S8C), consistent with prior findings (Pereira et al., 2015). All neurons identified through this dual-reporter approach are summarized in Figure S7C.

We compared our findings with previous reports and have compiled a list of potential co-transmitter neurons that are consistently identified across independent studies (Figure S9), suggesting these are likely *bona fide* co-transmitter neurons. Together, we observe that *C. elegans* has 35 neurons exhibiting molecular signatures of co-transmission (Table 2)—representing ~ 10% of the *C. elegans* nervous system (Figure S9 and Table 3). Strikingly, the pharyngeal nervous system—analogous to the vertebrate enteric nervous system (Albertson & Thomson, 1976)—had the highest density of co-transmitter neurons: 30% (6 of 20 neurons) displayed co-expression of multiple vesicular transporters. Across the entire nervous system, co-transmission was prevalent among sensory neurons, with 12% (10 of 83), compared to 11% of interneurons (9 of 81) and 7% of motor neurons (8 of 116) (Table 2). We also compiled a list of neurons that were suggested to be co-transmitters (RIB, AVA, AVB) (Gendrel et al., 2016), but that are not supported by other expression studies (Taylor et al., 2021) (Table 4).

**Table 2 –.** List of potential co-transmitter neurons by neuronal type.

| pharyngeal neurons | sensory neurons | interneurons | motor neurons |
| --- | --- | --- | --- |
| I2L | ADFL | AIML | HSNL |
| I5 | ADFR | AIMR | HSNR |
| M4 | AFDL | RIH | SMDDL |
| M5 | AFDR | PVNL | SMDDR |
| MI | URXL | PVNR | SMDVL |
|  | URXR | RIML | SMDVR |
|  | SDQR | RIMR | VC04 |
|  | DVA | RICL | VC05 |
|  | PDEL | RICM |  |
|  | PDER |  |  |

**Table 3 –.** List of potential co-transmitter neurons reported by at least more than one study.

| Co-transmitter neurons | Suggested by CenGEN expression data?* | Consistent with previous reports: | Combination suggested |
| --- | --- | --- | --- |
| **ADF** | Yes | Pereira et al. (2015), Wang et al. (2024) & this study | Serotonin + acetylcholine |
| **AFD** | Yes | Wang et al. (2024) & this study | Acetylcholine + glutamate |
| **AIM** | No | Jafari et al. (2011), Serrano-Saiz et al. (2013), Wang et al. (2024) | Serotonin + glutamate |
| **DVA** | Yes | Wang et al. (2024) & this study | Acetylcholine + glutamate |
| **HSN** | Yes | Duerr et al. (2001), Wang et al. (2024) & this study | Serotonin + acetylcholine |
| **I2L** | No | Wang et al. (2024) & this study | glutamate + UNC-47-dependent NT |
| **I5** | No | Jafari et al. (2011), Serrano-Saiz et al. (2013), Wang et al. (2024) | Serotonin + glutamate |
| **M4** | No | Wang et al. (2024) & this study | acetylcholine + UNC-47-dependent NT |
| **M5** | Yes | Wang et al. (2024) & this study | Acetylcholine + glutamate |
| **MI** | Yes | Serrano-Saiz et al. (2013), Wang et al. (2024) | Serotonin + glutamate |
| **PDE** | No | Wang et al. (2024) & this study | dopamine + glutamate |
| **PVN** | Yes | Wang et al. (2024) & this study | Acetylcholine + glutamate |
| RIC | Yes | Reilly et al. (2022), Wang et al. (2024) | glutamate + octopamine |
| RIH | Yes | Jafari et al. (2011), Wang et al. (2024) | Serotonin + acetylcholine |
| RIM | Yes | Alkema et al. (2005), Serrano-Saiz et al. (2013), Wang et al. (2024) | Glutamate + tyramine |
| **SDQR** | Yes | Wang et al. (2024) & this study | acetylcholine + UNC-47-dependent NT |
| **SMD** | Yes | Gendrel et al. (2016) & this study | GABA + acetylcholine |
| URX | No | Jafari et al. (2011), Wang et al. (2024) | Serotonin + acetylcholine |
| **VC04** | Yes | Duerr et al. (2001), Wang et al. (2024) & this study | Serotonin + acetylcholine |
| **VC05** | Yes | Duerr et al. (2001), Wang et al. (2024) & this study | Serotonin + acetylcholine |

*\*Determined with threshold of 1*

**Table 4 –.**
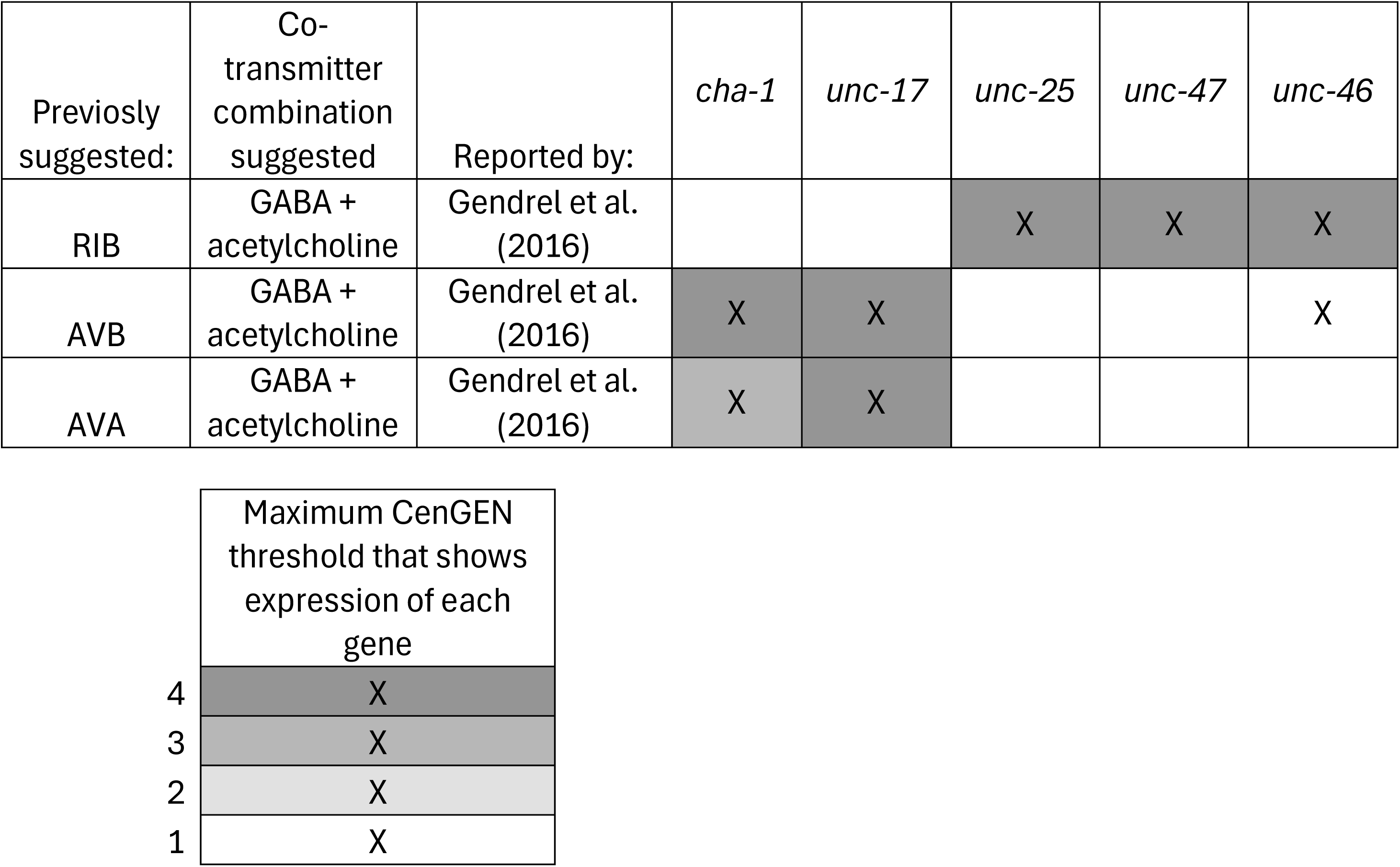
List of neurons inconsistent across studies to suggest co-transmission capacity.

| Previously suggested: | Co-transmitter combination suggested | Reported by: | *cha-1* | *unc-17* | *unc-25* | *unc-47* | *unc-46* |
| --- | --- | --- | --- | --- | --- | --- | --- |
| RIB | GABA + acetylcholine | Gendrel et al. (2016) |  |  | X | X | X |
| AVB | GABA + acetylcholine | Gendrel et al. (2016) | X | X |  |  | X |
| AVA | GABA + acetylcholine | Gendrel et al. (2016) | X | X |  |  |  |

|  |  |
| --- | --- |
|  | Maximum CenGEN threshold that shows expression of each gene |
| 4 | X |
| 3 | X |
| 2 | X |
| 1 | X |

### *In-vivo* visualization of co-transmitter synapses in the ADF chemosensory neuron

We next used our toolkit to investigate the subcellular localization of vesicular transporters in a co-transmitting neuron, ADF. The ADF neurons are a bilaterally symmetric pair of sensory neurons in *C. elegans* known to regulate food exploration, chemotaxis and entry into the lethargic-like dauer state (Bargmann & Horvitz, 1991). While ADF has long been known to use serotonergic neurotransmission, our findings indicate that it is also capable of acetylcholine synthesis and packaging (Figure 6H, S7C and S10A-B). Our findings are consistent with recent transcriptomic and reporter-based studies (Pereira et al., 2015; Taylor et al., 2021; Wang et al., 2024).

To examine the subcellular distribution of serotonin or acetylcholine vesicular transporters in ADF, we used previously developed tools to endogenously label the serotonin vesicular transporter CAT-1/VMAT (Huang et al., 2023), and the acetylcholine vesicular transporter UNC-17/VAChT with GFP (Figures 4 and 5). To label these vesicular transporters specifically in ADF, we drove Flippase and GFP1-10 expression using the AFD-specific promoter, *srh-142* promoter (Maicas et al., 2021). Both UNC-17/VAChT and CAT-1/VMAT displayed punctate fluorescence along the ADF axon, consistent with the location of presynaptic sites expected from electron microscopy-based 3D reconstructions (Figure S10C) (Koonce et al., 2024; White et al., 1986).

To then visualize the spatial relationship between these two vesicular transporters, we generated a strain in which UNC-17/VAChT was tagged with mRuby and CAT-1/VMAT with GFP at their respective endogenous loci, cell-specifically in ADF neurons (Figure 7A). *In vivo i*maging using a spinning disk confocal microscope revealed that these transporters frequently co-localize within the same synaptic boutons (Figure 7B and 7B’). Interestingly, we also found instances in which the UNC-17/VAChT::mRuby and CAT-1/VMAT::GFP signals partially segregate into separate boutons along the same axon (Figure 7C and 7C’), suggesting that these vesicular transporters can be sorted into distinct vesicle populations. Consistent with the idea that UNC-17 and CAT-1 localizes to synaptic vesicles along the axon, we observed that UNC-17 and CAT-1 localization was disrupted in a Kif1A/*unc-104* (*e1265*) mutant animal (Hall & Hedgecock, 1991), absent in the axon and accumulated in the cell body (Figure 7D-D’).

**Figure 7 –.**
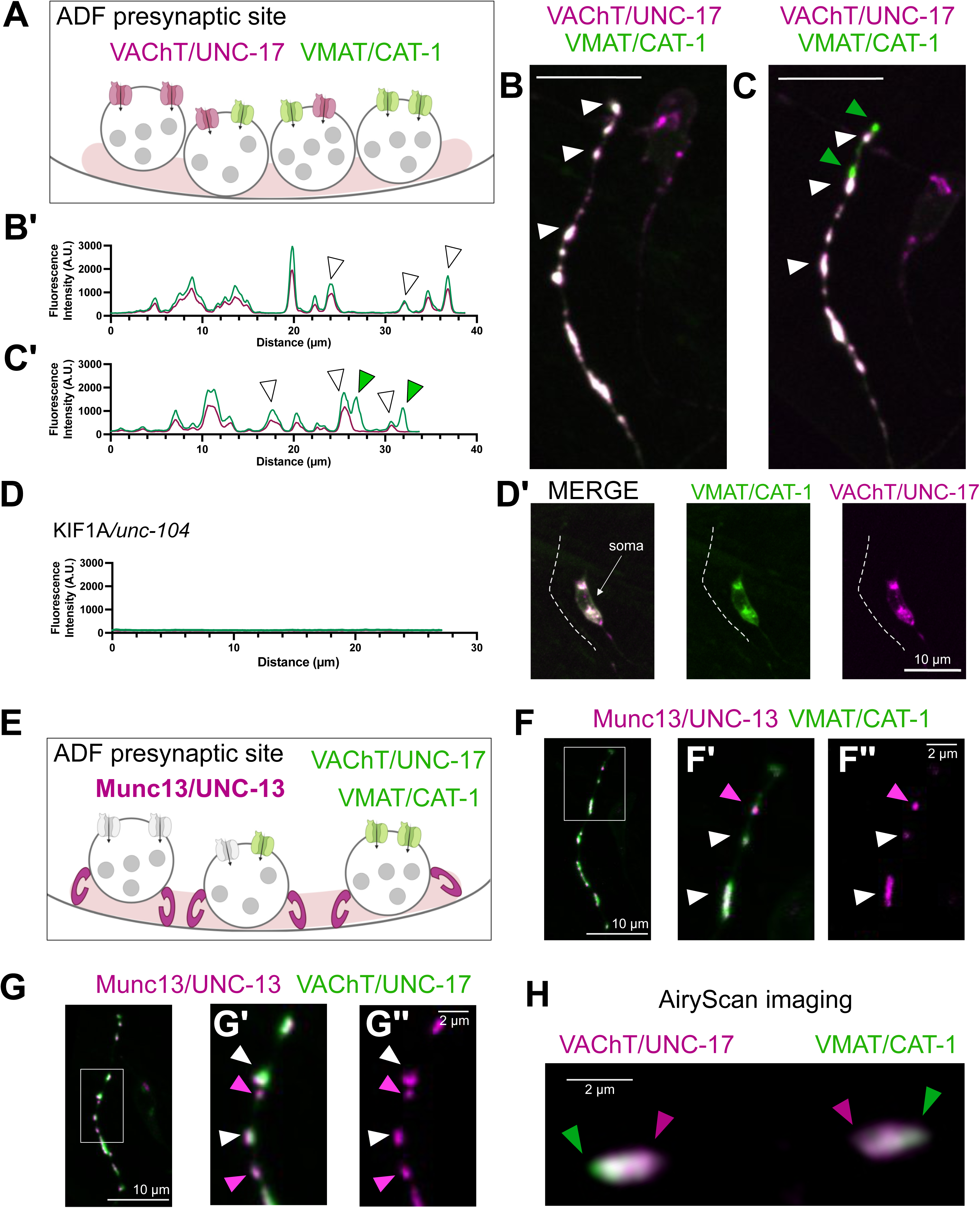
Visualizing the distribution of cholinergic and serotonergic vesicles in ADF neurons. **(A)** Dual-labeling of the endogenous acetylcholine (UNC-17, magenta) and serotonin (CAT-1, green) vesicular transporters in ADF neurons. **(B)** UNC-17::mRuby and CAT-1::GFP overlap along the ADF axon. White arrowheads denote overlap of both signals. **(B’)** UNC-17::mRuby and CAT-1::GFP intensity profile. **(C)** Example of UNC-17::mRuby and CAT-1::GFP when they partially do not overlap along the ADF axon, green arrowheads point to CAT-1-only puncta. **(C’)** UNC-17::mRuby and CAT-1::GFP intensity profile. Scale bar = 10μm. **(D-D’)** Endogenous UNC-17::mRuby and CAT-1:::GFP are absent from the axon of *unc-104* mutant animals. Line scan of axon **(D)** and fluorescent images **(D’)**. **(E)** Schematic of the dual-labeling of endogenous CAT-1::GFP or UNC-17::GFP with endogenous active zone protein UNC-13::mScarlet along the ADF axon. **(F)** Live imaging of the ADF axon reveals UNC-13::mScarlet puncta that lack CAT-1::GFP. Scale bar = 10 μm. **(F’-F’’)** Zoom-in region of E. White arrowheads show overlapping vesicular transporter tagged (green) with the active zone protein UNC-13 (magenta) in individual puncta. Magenta arrowhead points to UNC-13::mScarlet puncta that do not overlap with CAT-1::GFP. **(G)** Live imaging of the ADF axon reveals UNC-13::mScarlet puncta that lack UNC-17::GFP. Scale bar = 10 μm. **(G’-G’’)** Zoom-in region of F. White arrowheads show overlapping vesicular transporter tagged (green) with the active zone protein UNC-13 (magenta) in individual puncta. Magenta arrowhead points to UNC-13::mScarlet puncta that do not overlap with CAT-1::GFP. Scale Bar = 2 μm. **(H)** AiryScan imaging of dual-labeled UNC-17::mRuby and CAT-1::GFP along the ADF axon show localization in the same synaptic bouton but with distinct enrichment areas. Green arrowhead head points to CAT-1 enrichment and magenta arrowhead points to UNC-17 enrichment. Scale Bar = 2 μm.

To test whether both transporters are present at all ADF synapses, we endogenously tagged the active zone protein UNC-13/Munc13 with mScarlet and examined its spatial relationship to UNC-17/VAChT::GFP and CAT-1/VMAT::GFP (Figure 7E). We observed active zone UNC-13::mScarlet puncta that lacked either CAT-1/VMAT::GFP (Figure 7F) or UNC-17/VAChT::GFP labeling (Figure 7G). These findings are consistent with the idea that, while these two vesicular transporters display co-localization in synaptic varicosities, they also independently localize to distinct subcellular compartments.

Synaptic varicosities in *C. elegans* can be within the diffraction limits of light microscopy, preventing differentiation of co-localizing vesicular populations. To better understand the relative distribution of these vesicular transporters, we next visualized ADF synapses with increased resolution, using AiryScan imaging, which can achieve differentiation of fluorophores up to 120 nm apart (Wu & Hammer, 2021). We observed that, even in synaptic boutons in which the vesicular transporters were observed to co-localize with traditional light microscopy methods, UNC-17/VAChT::mRuby and CAT-1/VMAT::GFP differentially segregated when imaged using AiryScan microscopy (Figure 7H).

Together these results suggest that acetylcholine and serotonin co-localize to synapses, but might be packaged into distinct vesicles with specific synaptic subcellular localization that is detectable upon super-resolution microscopy. Our findings underscore the importance of endogenous labeling in determining the specific localization of these vesicular transporters and their use with higher-resolution imaging methods, highlighting the value of the tools developed in this study to understand the cell-biological organization of synapses *in vivo*, particularly for neurons using more than one neurotransmitter.

## DISCUSSION

The integration of anatomical connectivity, molecular identity, neural activity, and transmitter usage provides a powerful framework for building models of neural circuit function. The *C. elegans* community has access to a complete connectome (White et al., 1986); the cellular identity of all neurons (Sulston & Horvitz, 1977; Sulston et al., 1983); whole-brain calcium imaging (Nguyen et al., 2016; Prevedel et al., 2014; Schrödel et al., 2013); single-cell transcriptomic profiles (Taylor et al., 2021); and a full neurotransmitter identity map for all neurons (Wang et al., 2024). These datasets have inspired models describing how specific circuits may give rise to behavior. However, validating these models *in vivo* requires tools that can precisely manipulate the molecular components of individual synaptic connections (Dag et al., 2023; Hawk et al., 2018; Kumar et al., 2024). SynaptoTagMe provides that missing capability (Table 1) for the neurotransmitter systems that cover approximately 90% of the *C. elegans* nervous system (GABA, glutamate, acetylcholine, and the monoamines). By enabling cell-specific labeling and conditional knockout of vesicular transporters, we can now directly test the contribution of individual neurotransmitters within defined circuits and link those changes to behavioral outcomes. Moreover, due to the evolutionary conservation of vesicular transporters, the *in vivo* validation of tagging strategies will help identify suitable labeling strategies for other organisms, providing a path toward comparative and cross-species studies of synaptic dynamics based on neurotransmitter identity.

Co-transmission is a conserved feature of neural systems across the animal kingdom (Granger et al., 2017; Lacin et al., 2019; Vaaga et al., 2014), but its preponderance *in vivo*, its regulation and its functional significance is still an area of active research. Using *in-vivo* reporters subject to endogenous regulation, and contrasting our results with previous studies, we determine that more than 10% of *C. elegans* neurons have co-transmission potential (Figure S9) (Tables 2 and 3). Our *in vivo* characterization of co-transmitting neurons confirm and expand findings reported for the latest neurotransmitter atlas of *C. elegans* (Wang et al., 2024) and yield three key insights. First, co-transmission occurs throughout the nervous system of *C. elegans*, including both the pharyngeal (enteric-like) and more central nervous systems, like the nerve ring and nerve cords (Table 2). Second, neurons can co-transmit multiple neurotransmitters in specific combinations that are conserved from nematodes to mammals (Figure 6 and S9) (Granger et al., 2017; Lacin et al., 2019; Trudeau & El Mestikawy., 2018; Vaaga et al., 2014; Wang et al., 2024). Importantly, the same neurons consistently exhibit co-transmission of the same neurotransmitter identities across individual animals, consistent with co-transmitter identity mapping to neuronal identity (Figure 6D-H). Third, co-transmission is part of every layer of a circuit, from sensory neurons to interneurons and motor neurons (Table 2). This is especially interesting in light of recent studies showing that co-transmission in sensory and motor circuits can be modulated by environmental cues such as stress (Bertuzzi et al., 2018; Li et al., 2024; Pocock & Hobert, 2010) or light-dark cycles (Chen et al., 2023; Maddaloni et al., 2024). With the tools developed here - based on endogenously tagged vesicular transporters – it is now possible to monitor the dynamic expression and subcellular distribution of specific vesicle populations *in vivo* and what molecular mechanisms drive those changes.

Our conclusions are supported by independent and convergent lines of evidence, and the toolkit developed here enables direct empirical interrogation of both the existence and functional relevance of co-transmission *in vivo*. While expression of vesicular transporters represents one line of evidence for co-transmission potential, it does not by itself establish functional co-release, which additionally depends on neurotransmitter biogenesis, release competence, and activity-dependent regulatory mechanisms. SynaptoTagMe now make it possible to systematically test these requirements.

We note that the current characterization of co-transmitting neurons might be an under-estimate of the total number of neurons which use co-transmission. For example, it has been proposed that additional neurotransmitters, like betaine, may function in the *C. elegans* nervous system (Wang et al., 2024). Accounting for neurons that express proteins capable of synthesizing or packaging betaine, the proportion of potential co-transmitter neurons may exceed 20% of the whole nervous system of *C. elegans*. Our characterization of co-transmission focused on the *C. elegans* adult hermaphrodite, and co-transmitting neuron identities could be developmentally regulated, or modulated based on prior experience. Consistent with this, it has been observed that the identity of co-transmitting neurons is different between males and hermaphrodites (Serrano-Saiz et al., 2017), underscoring the importance of future examination of the plasticity and developmental regulation (Pereira et al., 2019; Pereira et al., 2015) of co-transmitting capacity for individual neurons.

Expression of a vesicular transporter, while consistent with co-transmitting capacity, is not conclusive for the existence of co-transmission for that neuron. For example, we identified co-expression of the GABA and Glutamate vesicular transporters in the pharyngeal neurons I2 (Figure S8A). Notably, I2 does not express the GABA synthesis enzyme, *unc-25*/GAD (McIntire, Jorgensen, & Horvitz, 1993) or the GABA re-uptake transporter, *snf-11* (Mullen et al., 2006). Thus, it is unlikely that it produces GABA or uptakes it from the extracellular space. VGAT/UNC-47 has also been reported to transport neurotransmitters such as glycine (Aubrey et al., 2007) and beta-alanine (Juge et al., 2013), raising the possibility that I2 could co-transmit glutamate with an unconventional neurotransmitter. Additionally, it is important to mention that the identification of *eat-4*-positive neurons through the replacement of the *eat-4* coding sequence and introns (Figure S7A) could result in the elimination of regulatory sequences. Thus, we conceptualize the list of co-transmitting neurons as a hypothesis-generating framework to be further examined with the tools developed in this study.

Our observations of the identity of co-transmitting neurons, and the specific combinations represented in the neurons, suggest that there may be transmitter-specific rules of synaptic biology important for circuit function (Silm et al., 2019; Trudeau & El Mestikawy., 2018). This is consistent with findings in vertebrates, in which specific neurotransmitter combinations and their distributions could underpin specific features of circuit function. For example, in Starburst Amacrine cells in the mammalian retina, acetylcholine and GABA (O’Malley et al., 1992) are packaged into distinct vesicle pools that exhibit different calcium sensitivities for release (Lee et al., 2010). The distribution of specific vesicular populations and their release probabilities might constitute an architecture that helps encode the sensory signals processed by Starburst Amacrine cells. We similarly hypothesize that the specific distribution of co-transmitting synapses across the *C. elegans* connectome, and the identities of the neurotransmitters used, might help encode features important for circuit function and animal behavior. By allowing longitudinal, cell-specific monitoring of the expression, regulation, and subsynaptic distribution of distinct vesicle populations in living animals, this toolkit provides a platform for probing when, where, and how co-transmission is deployed, and for defining the molecular and circuit-level mechanisms that govern its use under different physiological and environmental conditions.

## METHODS

### Strains

Worms were maintained at 20°C using standard techniques (Brenner, 1973). Strains were maintained on NGM plates seeded with *E. coli* (OP-50). The wild type (WT) is N2, and only hermaphrodite worms were used for this study. A complete list of strains appears below. Strains that were developed in this study as part of the SynaptoTagMe toolkit and appear in Table 1 can be requested through CGC. The necessary sequencing information can be found at: https://www.intralab.app/research-papers/cuentas-condori_etal-2026.

### Generation of new alleles

For the strains engineered by Sunybiotech, as described below, strain design was performed in the Colón-Ramos lab by Andrea Cuentas-Condori.

Sunybiotech used CRISPR/Cas9 to insert GFP FLP-on cassettes (Schwartz & Jorgensen, 2016) at either the N-termini of the *unc-17* locus (*syb7251*) or C-termini end of the *unc-17 (ola503),* or *eat-4 (syb8568)* locus, according to sequence design. mRuby FLP-on cassettes were inserted similarly at the C-termini end of *eat-4* (*syb9193*) and *unc-17* (*syb7882*) locus. To visualize the fluorescent signal tagged to the protein of interest, Sunybiotech generated single-copy MosCI strains.

Sunybiotech used CRISPR/Cas9 to add full length GFP (*syb6990*), full length mKate (*syb7358*) or split GFP (one (*syb7313*) or three (*syb7849*) copies of GFP11) to the *unc-47* locus at the +893bp position. To visualize the reconstituted GFP signal in RIB neurons, complementary GFP1-10 was driven with the P*sto-3b* promoter; and to visualize the GFP-reconstituted signal in DD neurons, GFP1-10 was driven with the P*flp-13* promoter.

Sunybiotech used CRISPR/Cas9 to introduce an FRT site before the +1bp in the *unc-25* gene locus (*syb5949*). In a second round of CRISPR editing, they introduced *let-858* 3’ UTR followed by a second FRT and nuclear mCherry (*syb6275)*.

*cat-1* (*ky1101 ky1118*) cell-specific knock out strain was created using CRISPR/Cas9 to introduce an FRT site immediately before the ATG in the *cat-1* gene locus (*ky1101*). A second round of CRISPR editing in that strain introduced the *let-858* 3’ UTR followed by a second FRT and the mCherry coding region immediately after the stop codon of *cat-1* to generate *ky1118*.

Sunybiotech used CRISPR/Cas9 to introduce a T2A::Flippase sequence before the STOP codon of *unc-47(syb8125)* and *unc-17(syb8059)*. All strains generated using CRISPR/Cas9 were outcrossed twice before use.

### Molecular Biology

Plasmids were constructed using Gibson cloning. First, Snapgene (Version 7.0.3) software was used to design primers targeting the desired DNA vector backbone and DNA insert. The vector backbone and DNA insert were PCR linearized and amplified using “CloneAmp HiFi PCR Premix.” To assemble the desired plasmid, the purified vector backbone DNA and insert DNA were combined and incubated in solution with “2x Gibson Assembly Enzyme Premix.” Following incubation, the reaction mixture was used to transform Stellar Competent Cells, which were subsequently plated and grown overnight on LB-Amp plates. All plasmids were verified with Sanger sequencing.

### Protein alignment and structure visualization

For each gene under study, NCBI BLAST was used to generate a protein sequence alignment of the *C. elegans* gene with the closest orthologs from the other model organisms *M. musculus*, *D. rerio*, *D. melanogaster*, and *H. sapiens*. Protein structure models for the C. elegans genes were downloaded from the AlphaFold database (Jumper et al., 2021) and predicted models for the CRISPR-Cas9 modified genes including fluorophores were generated by the Alphafold3 online server (Abramson et al., 2024). Visualization and image generation of protein structures was done using the ChimeraX software (Pettersen et al., 2021). To color the structures by sequence conservation, the alignments per gene were overlaid onto the structures with ChimeraX and colored by resulting sequence conservation Z-scores as calculated by the AL2CO algorithm (Pei & Grishin, 2001) within the software.

**Table 5 –.** List of all molecular locations tested for labeling each synaptic vesicle transporter.

| Gene | AA tested for fluorescent protein insertion | Functional tag? |
| --- | --- | --- |
| *eat-4* | N-terminus (After START codon) | yes |
| *unc-47* | Between E145 – N146 | yes |
| *unc-17* | Between AA P6-V7 | yes |
| *unc-17* | Between AA N242 - P243 | no |
| *unc-17* | C-terminus (Before STOP codon) | yes |

### Microscopy

Larval or young adult animals were immobilized on 2-10% agarose pads with 10mM levamisole. A Nikon Ti2 microscope equipped with a CSU-W1 spinning disk head, ORCA-Fusion BT SCMOS camera, high-speed piezo stage motor, 60X/1.40 Apo Lambda oil objective lens was used for live imaging. Z-stack images were collected (0.3-0.5 μm/step), spanning the focal depth of the nerve cord and nerve ring synapses. A Zeiss LSM880 microscope equipped with an AiryScan detector and 63X NA 1.4 oil objective was used for AiryScan imaging. FIJI (Schindelin et al., 2012) or NIS Elements AR analysis software (version 6.10.01) were used to create maximum intensity projections and 2D renderings.

### Thrashing Assay

*C. elegans* were raised at 20°C under standard laboratory conditions on agar plates seeded with a lawn of *E. coli* (OP50). Worms were synchronously grown to L4-stage and placed in individual wells of a Corning™ PYREX™ Spot Plate (Catalogue #722085) containing 1000 μl of M9 buffer, ensuring the buffer remained within the well’s borders. After a 30-second acclimation period to M9, thrashes were manually counted for 1 min. A single thrash was defined as a change in the direction of the worm’s midbody bending, counting each time the worm’s body flexed to one side. Following each trial, the worm was removed using a pipette and disposed of, and the M9 buffer was absorbed and discarded. The well was then cleaned with 70% ethanol and wiped dry. To avoid bias, the counter was blinded to each genotype. Each worm was tested only once, with assays conducted on 10 worms per genotype per day, and repeated over 2–3 days to account for potential day-to-day environmental variations.

### Chemotaxis Assay

Worms were maintained at 20°C for at least two generations on Nematode Growth Medium (NGM) seeded with OP50 *Escherichia coli* bacteria. The concentration of our attractant (NaCl) is approximately 50 mM in NGM plates. “Training” plates were produced using NGM with the further addition of 50 mM NaCl to a total concentration of 100 mM, then also seeded with OP50.

Chemotaxis assay was modified from standard procedure (Ward, 1973). All assays were performed on 50 mm diameter plates. Unseeded NGM plates were marked at the center and one point 12.5 mm away from the center. A ~60-85 mM gradient of NaCl was created between the center and outer point by adding 5 M NaCl at the outer point as drops of 4μL (20-24 hours before the assay), 4μL (5 hours before), and 1.6μL (2 hours before); a sham gradient was created using only water. Gradient prediction was determined as previously described (Crank, 1956; Pierce-Shimomura et al., 1999); briefly, for every point some distance *r* in cm from the salt peak, the concentration *C* in mM at any point in time was calculated as:

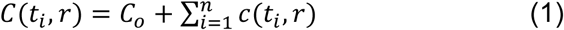

where *C_o_* is the initial concentration of NaCl in the agar (50 mM), *n* is the drop number, and *t_i_* is the time in seconds since the drop had been applied; the contribution from each drop, in turn, was calculated as:

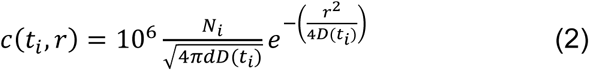

where *N_i_* is the moles of NaCl added per drop; *d* is the depth (cm) of the agar; and *D* is 1.590 × 10^-5^ cm^2^/s, the diffusion coefficient for 5 M NaCl through an aqueous medium (Robinson RA, 1959). The resulting gradients were validated by electrical conductivity measurements using an Oakton CON 6+ Handheld Conductivity Meter with a custom conductivity probe with 1 mm insertion depth (Micro-electrodes, Inc, Bedford, NH). The conductivity readings from 50 mM and 100 mM NaCl NGM plates were used for calibration at specific room temperatures.

The day before experiments, L4 animals were transferred to a seeded NGM plate to synchronize worms by developmental stage. At 5 hours before each assay, worms were transferred to a training plate using standard NGM plate recipe adjusted to 100 mM NaCl. After training, 8 worms were picked, with preference to those on the bacterial lawn, washed sequentially in two 100 µL drops of liquid NGM buffer (25 mM potassium phosphate pH 6, 1 mM CaCl2, 1 mM MgSO4, 50 mM NaCl) to remove adherent OP50, and transferred to a single 2 μL drop of NGM buffer at the center of the assay plate with prepared NaCl gradient as described. Data collection began when the water drop was fully absorbed into the assay plate and the first worm began to migrate from its starting point. Six assay plates were imaged for each strain across two separate days, yielding a total of 48 worms imaged per strain.

Images of chemotaxis behavior were acquired at 3.75 fps for 7 minutes using a Basler acA2440-35mm monochromatic sensor with an infrared filter on a commercially available WormLab imaging system and computer running WormLab 2023.1.1 software (MBF Bioscience LLC, Williston, VT USA). Individual worm position data was obtained by constructing tracks in WormLab software, then analyzed using custom scripts in R 4.4.1 (can be accessed through GitHub (https://github.com/colonramoslab/Cuentas-Condori-et-al.-2025-Toolkit-). The assay outcome was defined as the mean distance from the peak of the salt gradient for each worm, averaged over every available frame in the last minute of the assay. When a worm track was interrupted, e.g. by a worm exiting the camera field of view or by two worms intersecting, the last available position for the worm was repeated until the worm was re-detected.

### Roaming Assay

Roaming assay plates were prepared 3-5 days prior to the experiment by seeding NGM agar plates with *E. coli* (OP50) culture no older than 2 days. Plates were seeded using a sterile glass rod to spread the bacteria evenly across each plate. Plates were left to dry completely between 3-5 days at room temperature to ensure the bacteria layer was fully dry, thereby allowing for visible worm tracks during the assay. *C. elegans* were raised at 20°C under standard laboratory conditions on agar plates seeded with a lawn of *E. coli* (OP50). On the day before the assay, worms synchronously grown to L4-stage were transferred to regular seeded plates and stored at 20°C. After approximately 10 hours, worms were transferred to individual assay plates and incubated at 20°C for 16 hours. After this time, worms were removed and a grid overlay (3 mm x 3 mm squares) covering the assay plate was used to count the number of squares the worms had traversed during the incubation period. The number of squares crossed provided a quantifiable measure of roaming activity. To avoid bias, the counter was blinded to each genotype. Each worm was tested only once, with assays conducted on 10 worms per genotype per day and repeated over 2–3 days to account for potential day-to-day environmental variations.

### Aversion behavior Assay

Aversion behavior assay were performed as previously described (Feng et al., 2025). Animals were fed on *E. coli* BW25113 for at least three generations before the behavioral assay. 12.5 μL of overnight BW25113 cultures were seeded onto standard NGM agar plates, grown at 37°C incubator for 24 hours and then left at room temperature for another 24 hours. 15~20 animals at L4 stages from each genotype were transferred onto behavioral assay plate and recorded at 21°C for 20 hours, at a recording rate of 1 frame per minute. Biological replicates across two different days were conducted. Videos were cropped and analyzed using standard MatLab codes (Marquina-Solis et al., 2024). Aversion ratio was defined by the number of worms outside the bacterial lawn over total number of worms on assay plates.

### Statistical Analysis

We used the Shapiro-Wilk test to determine sample distribution. For comparisons between 2 normally distributed groups, Student’s T-test was used and p<0.05 was considered significant. ANOVA was used to compare between 3 or more normally distributed groups followed by Dunnett’s multiple-comparison test. If the samples were not normally distributed, we used a Mann-Whitney test to compare two groups and a Kruskal-Wallis test to compare three or more groups. Specific post-hoc statistical tests are listed in the figure legend of each experiment. Prism 10.4.2 was used to graph the data and for all statistical analysis.

### List of strains

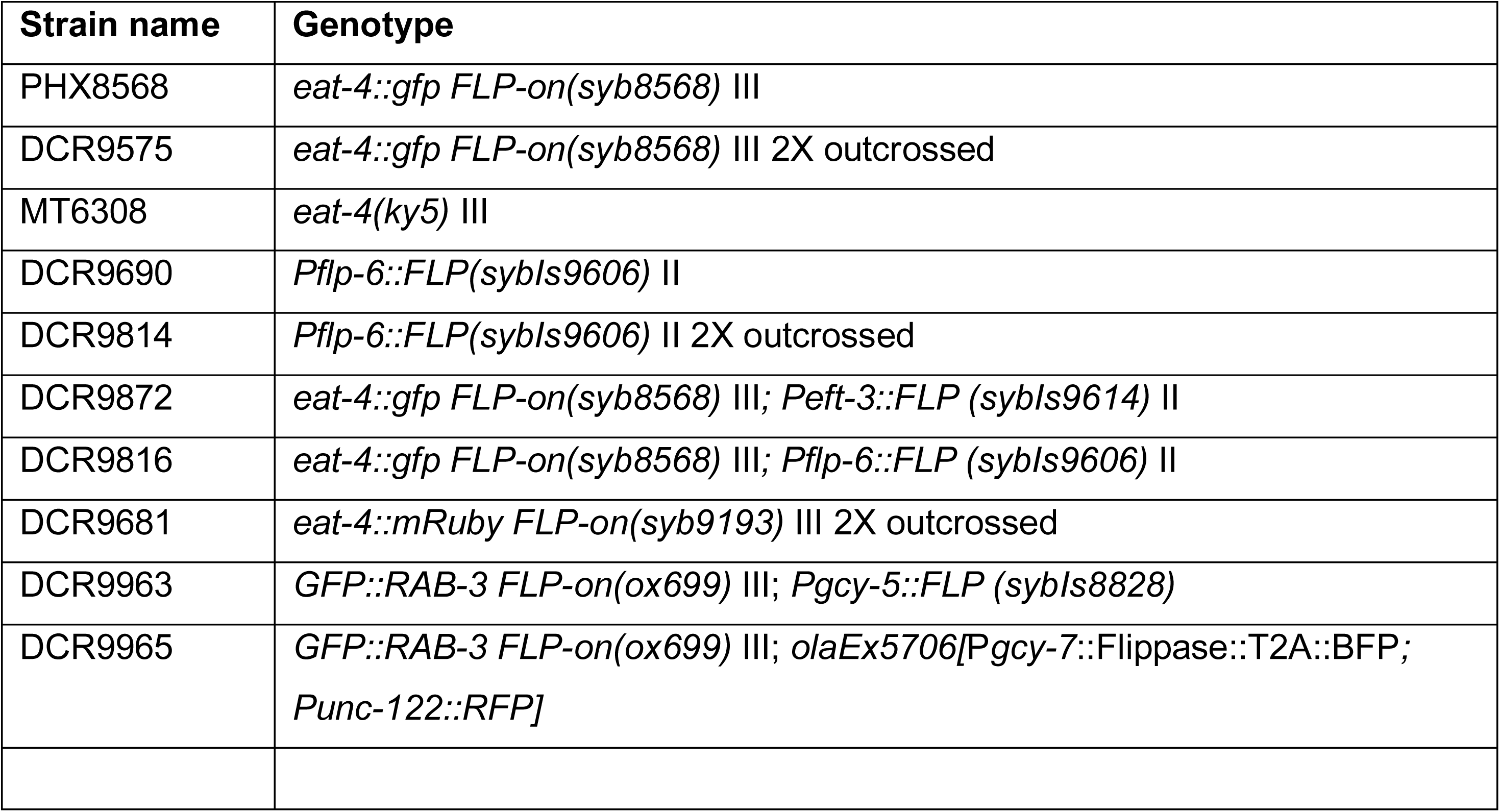
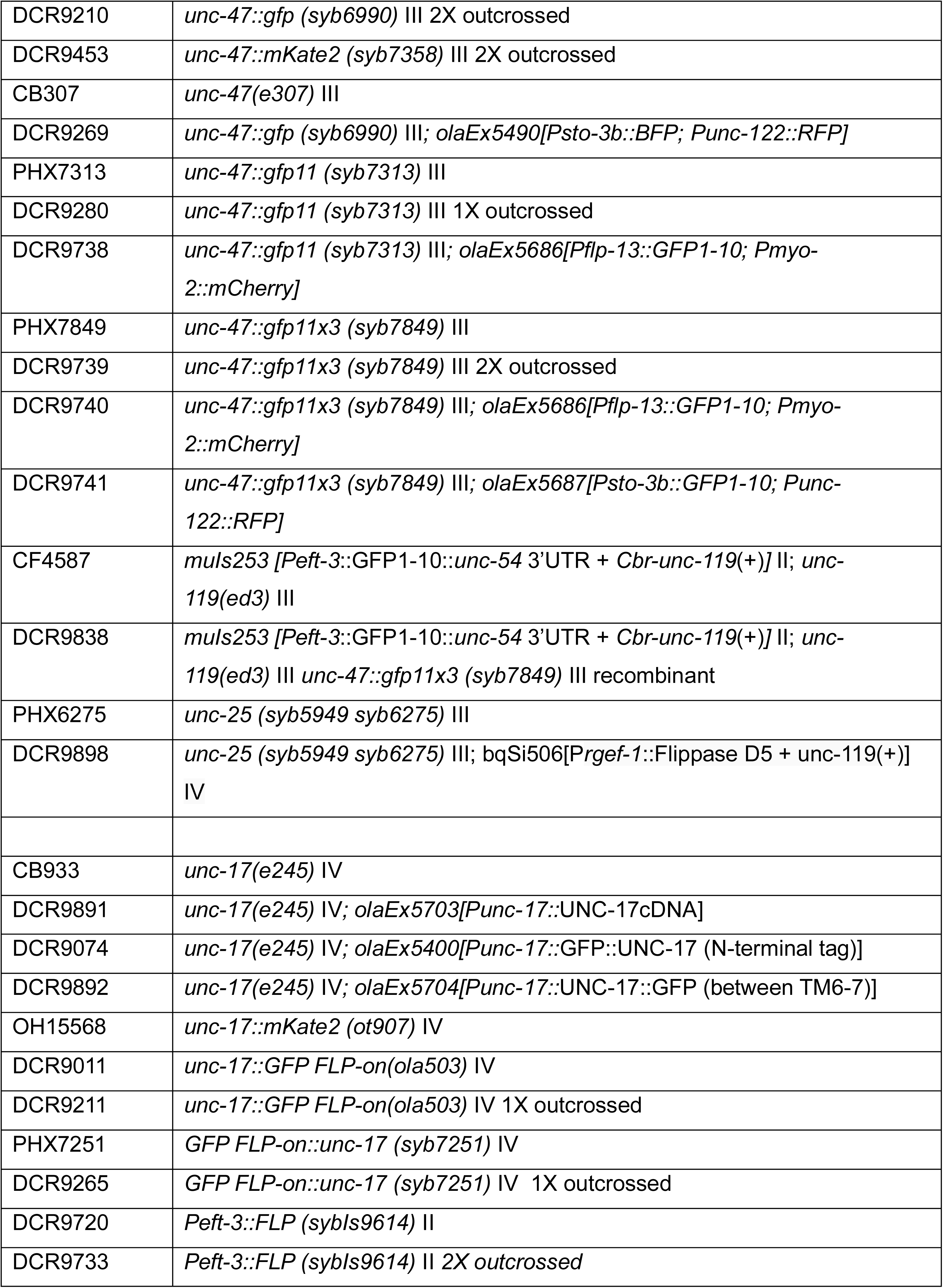
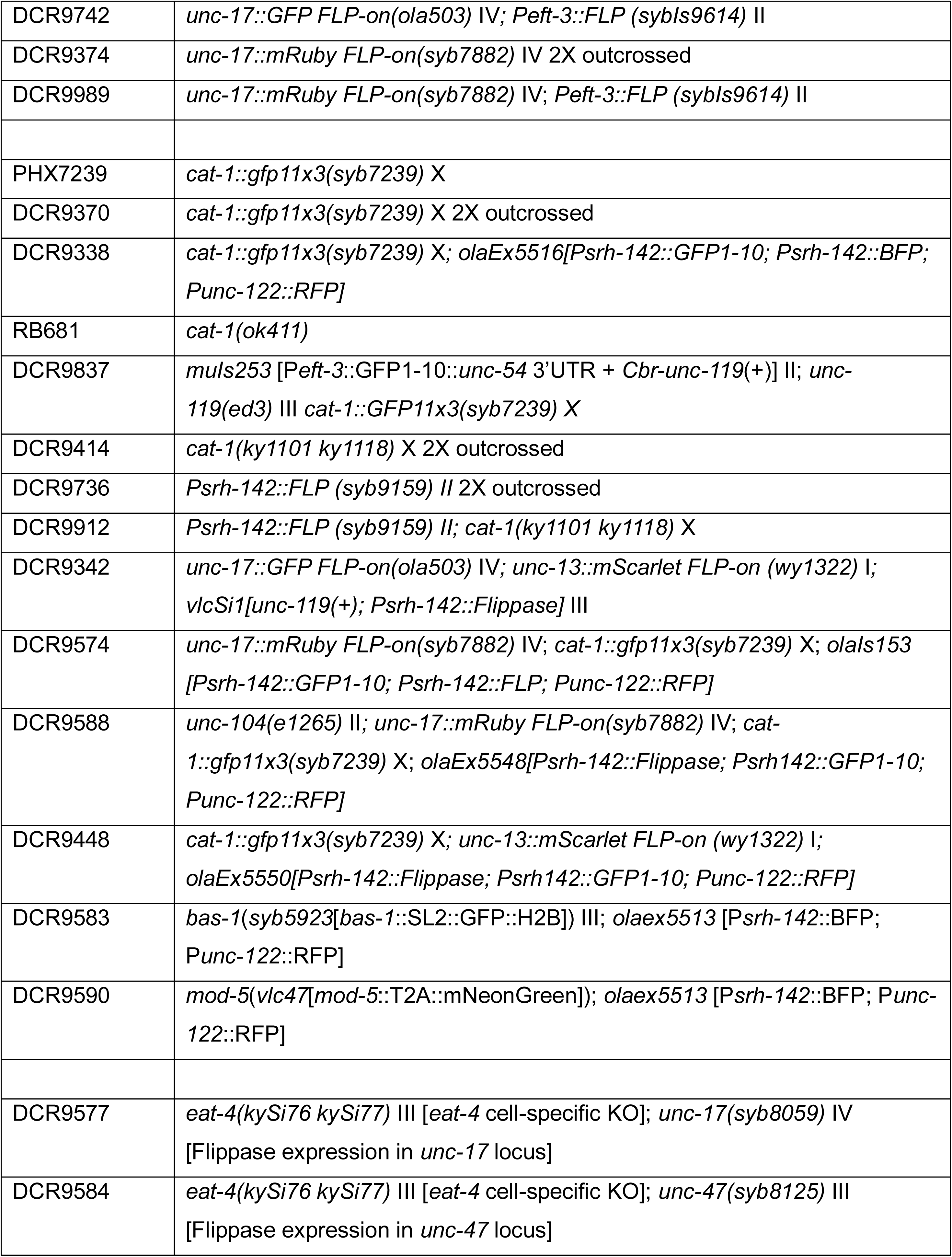
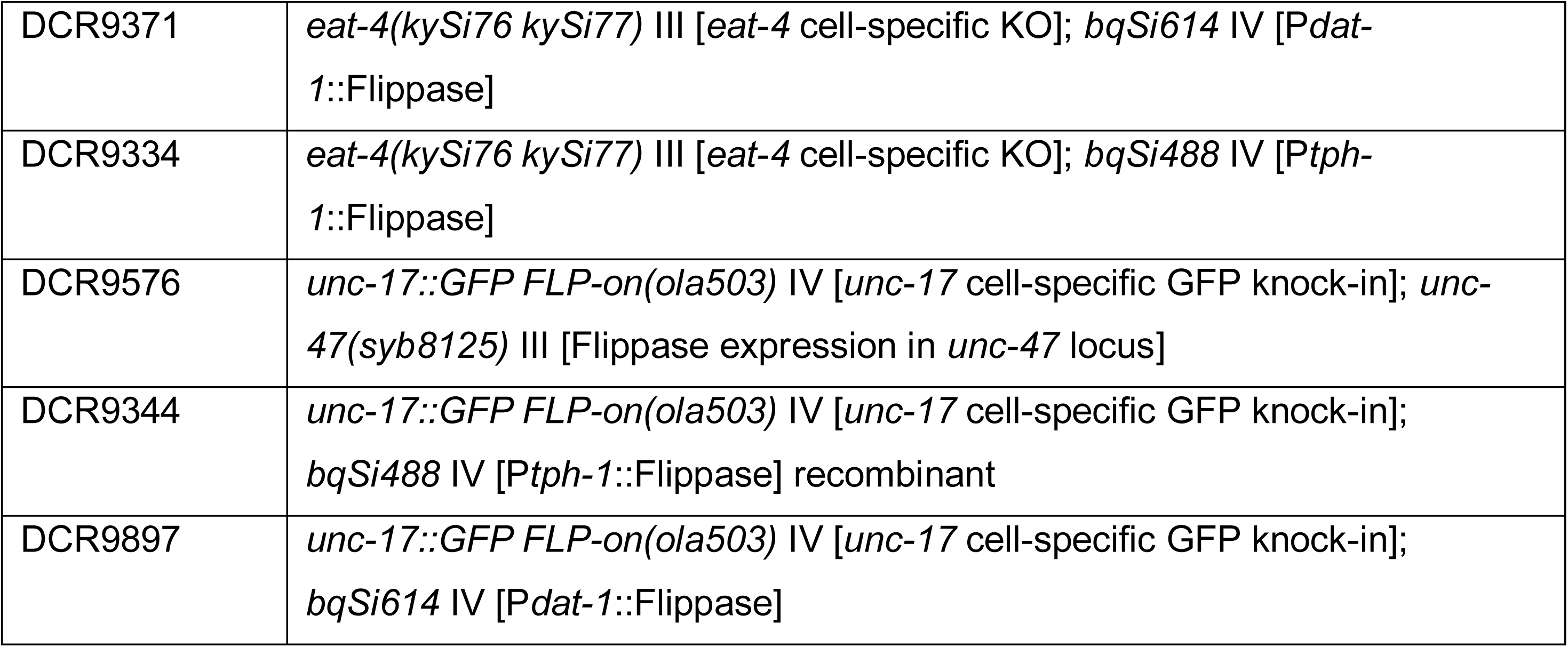

### List of plasmids

| Plasmid name | Genotype |
| --- | --- |
| DACR218 | *Punc-122::RFP* |
| DACR704 | *Pmyo-2::mCherry* |
| DACR4016 | *Psto-3b::BFP* |
| DACR4027 | *Punc-17::UNC-17cDNA* |
| DACR4033 | *Punc-17::UNC-17cDNA::GFP* in TM6-7 |
| DACR4038 | *Punc-17::GFP::UNC-17cDNA* |
| DACR4064 | *Psto-3b::GFP1-10* |
| DACR4078 | *Psrh-142::BFP* |
| DACR4083 | *Psrh-142::Flippase* |
| DCR4092 | *Psrh-142::GFP1-10* |
| pSH87 | *Pflp-13::GFP1-10* |

## ACKNOWLEDGEMENTS

We thank Nuria Flames Bonilla (Instituto de Biomedicina de Valencia, Spain), Rafa Alis (Instituto de Biomedicina de Valencia, Spain), Steven Flavell (MIT), and Yung-Chi Huang (MIT) for sharing reagents and constructs. We thank James Rand, Oliver Hobert (Columbia University) and Chen Wang (Columbia University) for sharing unpublished observations. We thank Ian Gonzalez for the creation of the name for this toolkit. We thank members of the Colón-Ramos Lab for feedback on figures and the manuscript. We also thank Stacy Wilson for technical support and training using the AiryScan imaging set-up as part of the Yale Neuroscience Imaging Core Facility. We also thank Emerson Santiago, member of the Koelle Lab (Yale University), for expert advice on assays of serotonin function. Some strains were provided by the CGC, which is funded by NIH Office of Research Infrastructure Programs (P40 OD010440). Some figures were created in https://BioRender.com. This work was supported by National Institutes of Health grants to DC-R (R35NS132156 and R01NS076558) and to AW (K99AG083129), to EMJ (R01 NS034307), and to MLS (F32GM133139). The Pew Latin American Fellowship to ACC (AWD0006561), the Jane Coffin Childs Fellowship to ACC (AWD0006564), the HHMI Hanna Gray Fellowship to ACC (GT15993), and the Chan Zuckerberg Initiative to CIB.

---

**Figure S1.**
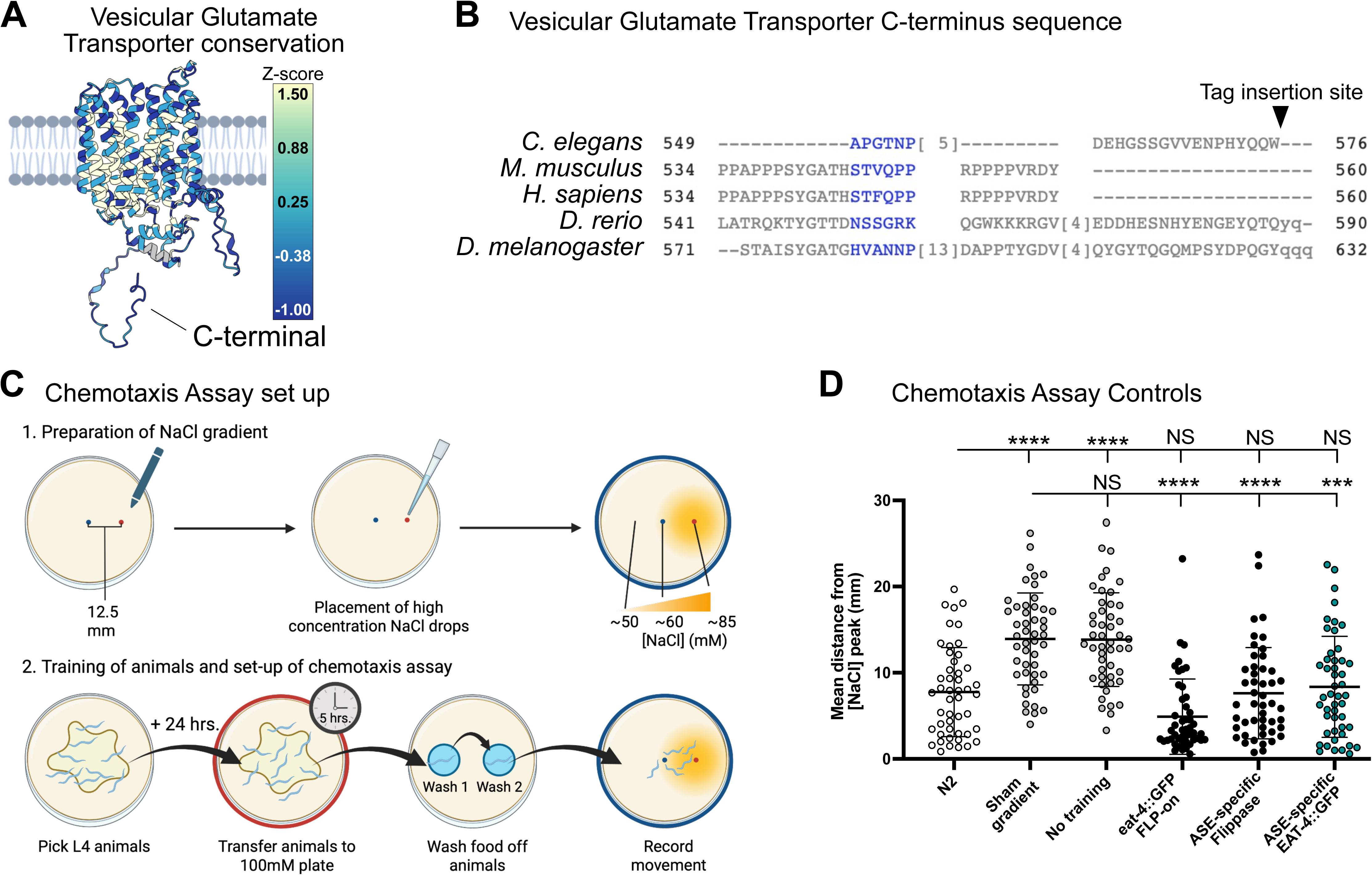
**(A)** Graphical representation of conservation of the amino acid predicted structure for the Vesicular Glutamate Transporter across common model organisms (named in Figure S1B). Dark colors denote the least conserved region, while clear colors denote the highest conservation. Note the C-terminus end, where we introduced the GFP, is one of the darkest (least conserved) regions in the structure. **(B)** Sequence alignment of the amino acid sequences for the Vesicular Glutamate Transporter across common model organisms. Blue letters indicate conservation of the amino acid properties (basic, acid, neutral) but not identities. Gray represents columns with gaps and no conservation. **(C)** Schematic of chemotaxis assay. (Top) A concentration gradient of NaCl (~60-85 mM) is established on assay plates (blue rim) by adding high concentration NaCl drops to the outer point of the assay plate at pre-determined times (See Methods). (Bottom) Larva stage 4 animals are picked, and twenty-four hours later, the young adult animals are transferred to NaCl 100mM training plates (red rim) for five hours. Animals are transferred to NGM buffer drops to wash off bacteria, before being placed on the assay plates (blue rim, placement at blue dot in schematic), where animal movement away from the highest [NaCl] is recorded (towards red dot in schematic). **(D)** Displacement of trained wild type animals on a sham gradient (13.92 ± 5.3 mm) or untrained wild-type animals on a gradient of NaCl (13.84 ± 5.4 mm). Wild-type animals (7.76 ± 5.2 mm) migrate across the salt gradient like EAT-4::GFP FLP-on (*syb8568*) animals that express (8.36 ± 5.8 mm) flippase in ASE neurons (*sybIs9606*). Animals that only express flippase in ASE neurons (7.61 ± 5.3 mm) migrate across the salt gradient similar to wild-type animals. Results represent the mean distance of each worm from the salt peak, averaged across the final minute of the assay, with each dot representing an individual animal. Plots are overlaid with Mean ± Standard Deviation. Kruskal-Wallis test with Dunn’s multiple comparison post hoc test. **** represents p<0.0001; *** represents p<0.001; * represents p<0.05; and NS means “not significant”.

**Figure S2.**
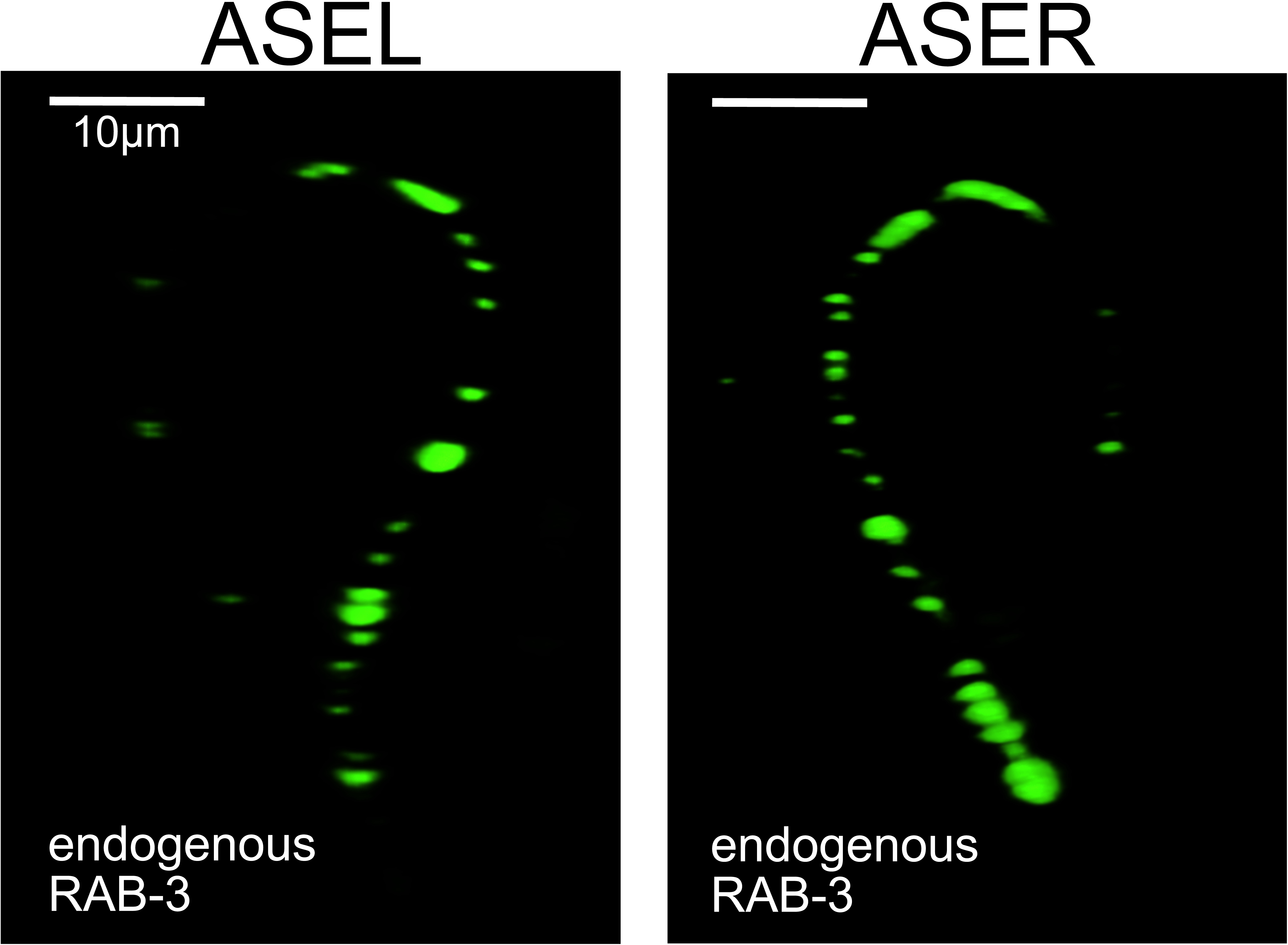
Endogenous RAB-3::GFP synaptic vesicle clusters in the axonal process of ASER and ASEL.

**Figure S3.**
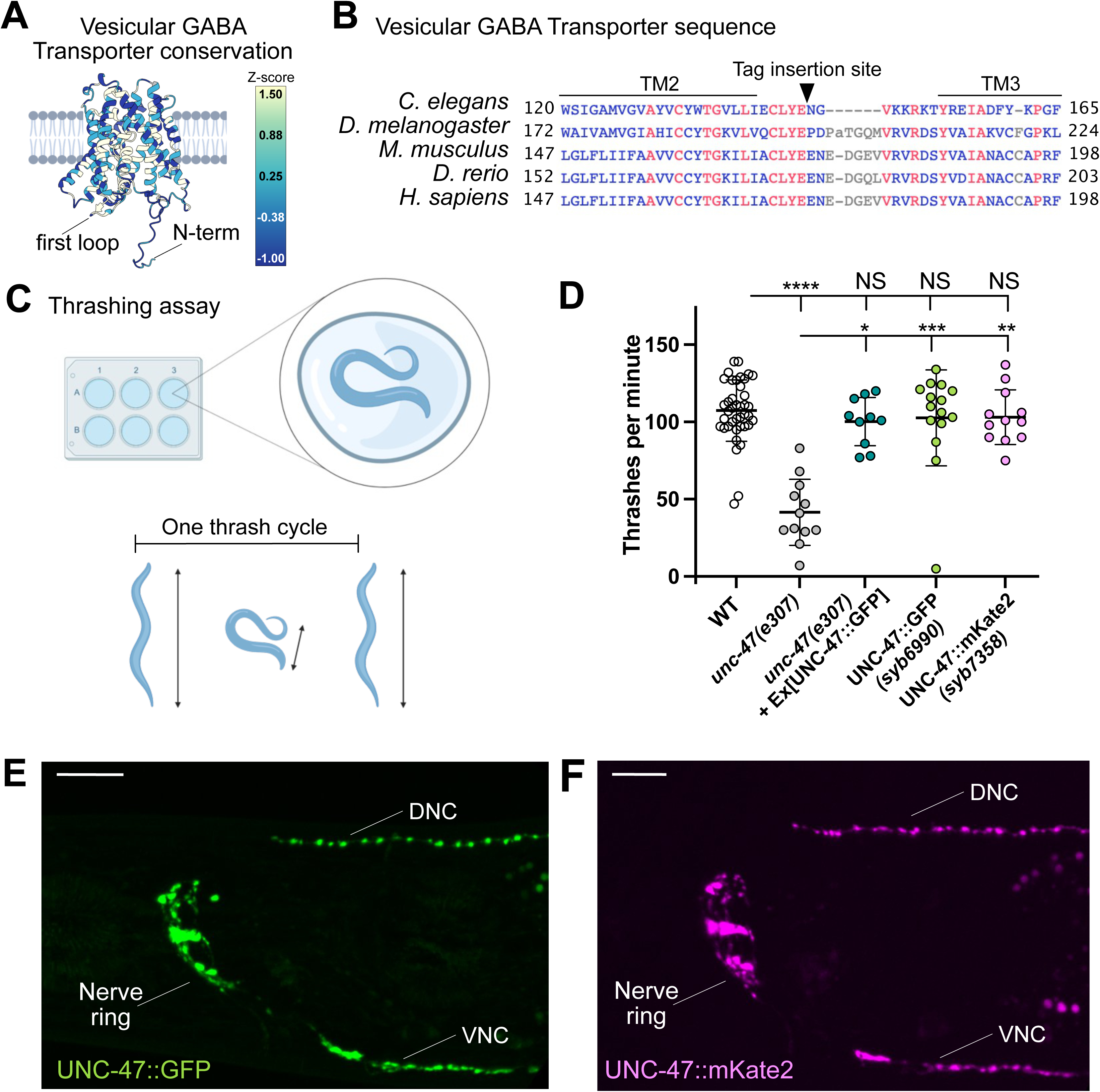
**(A)** Graphical representation of conservation of the Vesicular GABA Transporter amino acid predicted structure across common model organisms (named in Figure S2B). Dark colors denote the least conserved region, while clear colors denote the highest conservation. **(B)** Sequence alignment of the cytosolic loop between transmembrane domains 2 and 3 of the vesicular GABA transporter across common model organisms. Red letters indicate highly conserved columns (conservation of amino acid identity), and blue letters indicate conservation of the amino acid properties (basic, acid, neutral) but not identities. Gray represents columns with gaps. **(C)** Schematic of thrashing assay. “One thrash cycle” is scored when the animal bends as indicated in the schematic (from elongated-to-bent-to-elongated). We measured the number of thrashes per minute. **(D)** *unc-47(e307)* mutant animals thrash significantly less (41.5 ± 21.35) than wild-type animals (109.2 ± 23), while over-expression of UNC-47::GFP (100.2 ± 16), CRISPR knock-in UNC-47::GFP (107.9 ± 29) and UNC-47::mKate2 (103.1 ± 18) CRISPR-tagged animals thrash similarly to wild-type animals. Plots are overlaid with Mean ± Standard Deviation. Kruskal-Wallis test with Dunn’s multiple comparison post hoc test. **** represents p<0.0001; *** represents p<0.001; ** represents p<0.01; * represents p<0.05; and NS means “not significant”. **(E-F)** Fluorescent image of endogenously tagged **(E)** UNC-47::GFP (*syb6990*) and **(F)** UNC-47::mKate2 (*syb7358*). DNC = Dorsal Nerve Cord, VNC = Ventral Nerve Cord. Scale Bar = 10μm.

**Figure S4.**
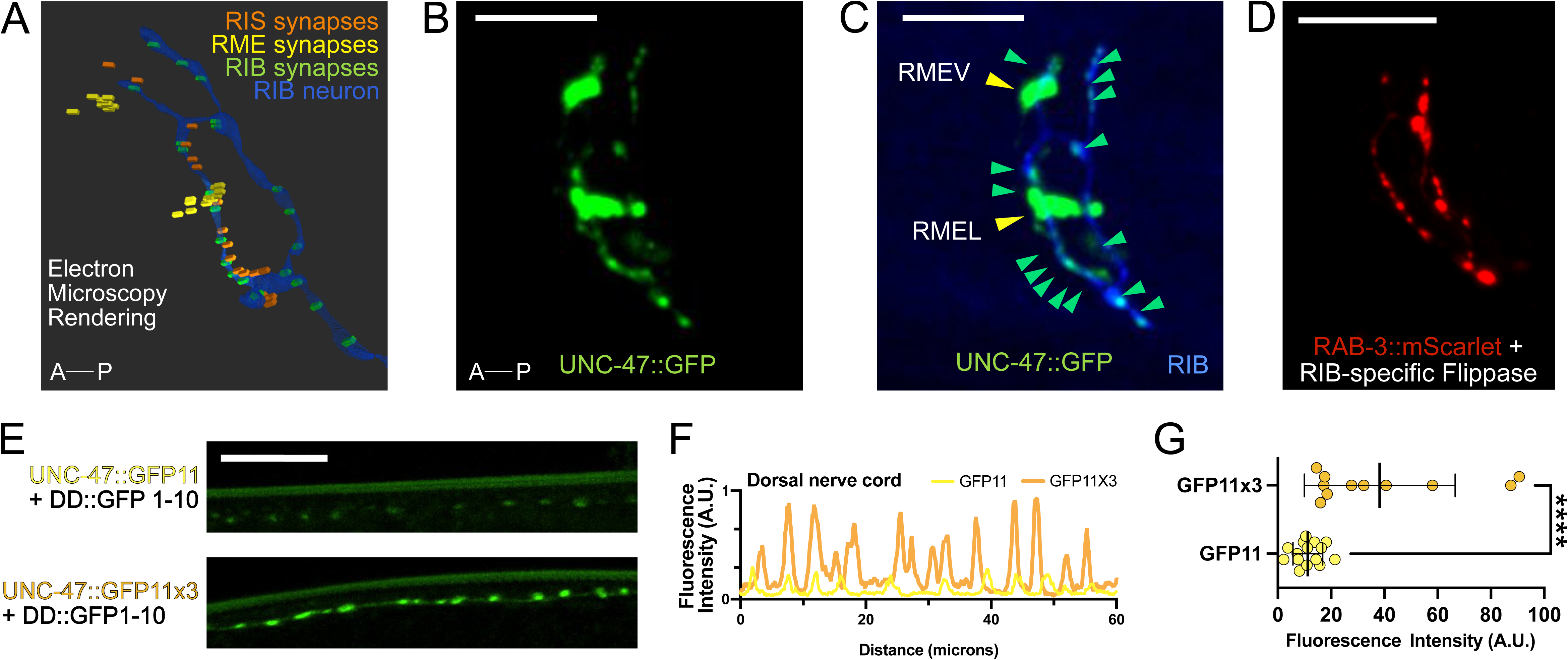
**(A)** All presynaptic sites from known GABAergic neurons in the *C. elegans* nerve ring (RIS, RME, and RIB), according to 3D electron microscopy of a Larva stage 4 (L4) wild-type animal (White et al., 1986) (image generated with NeuroSC (Koonce et al., 2025)). **(B)** 3D-rendering of UNC-47::GFP (green) puncta in the *C. elegans* nerve ring. A-P denotes anterior-posterior axis. Scale bar = 10 μm. **(C)** Addition of RIB::BFP into Figure S3B. Green arrowheads point to UNC-47::GFP puncta that overlap with RIB::BFP. Yellow arrowheads point to the synapses we interpret, based on our labeling and distribution, to belong to the RME neurons. **(D)** Endogenous mScarlet::RAB-3 in RIB highlight the characteristic shape of the RIB axon. **(E)** *In-vivo* reconstitution of UNC-47::GFP11x3 in DD neurons when tagged with (Top) one or (Bottom) three copies of GFP11 (P*flp-13*::GFP1-10) (He et al., 2019). **(F-G) (F)** Line scans and **(G)** quantification of reconstituted GFP fluorescence intensity with one (11.3 ± 6) or three copies (38.3 ± 28) of GFP11 in DD neurons. Scale bar = 10 μm. Plots are overlaid with Mean ± Standard Deviation. Mann-Whitney test. **** represents p<0.0001.

**Figure S5.**
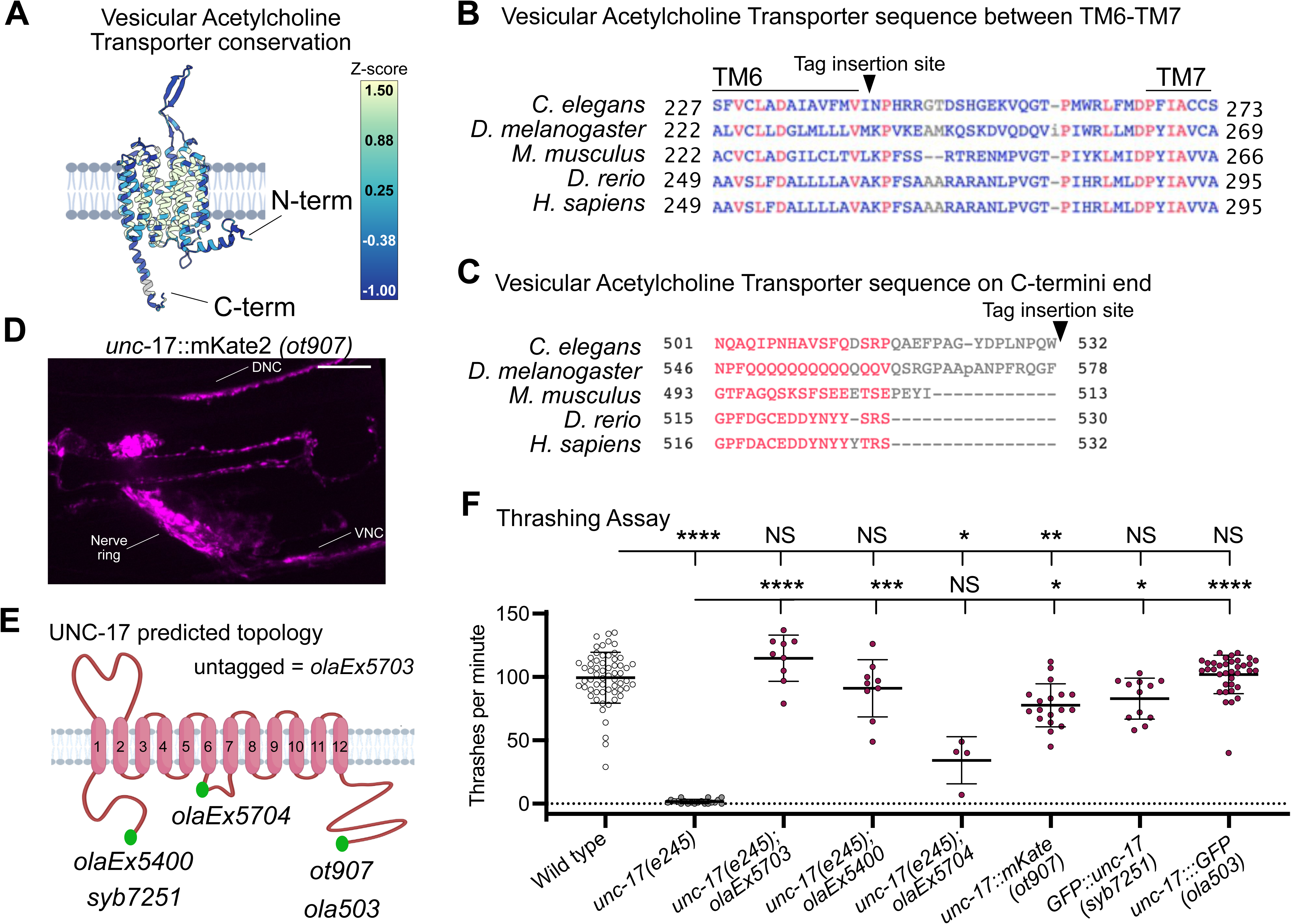
**(A)** Graphical representation of the amino acid conservation of the Vesicular Acetylcholine Transporter (VAChT) across common model organisms. Dark colors denote the least conserved region, while clear colors denote the highest conservation. Note the C-terminus end is one of the darkest regions in the structure, where the fluorophore was added. **(B-C)** Amino acid sequence alignment of VAChT across model organisms. Black arrowhead points to insertion site of fluorescent tag. **(B)** Sequences between transmembrane domains 6 and 7, which was tagged in *olaEx5704* array (See panel E). **(C)** Sequence at the C-terminus end. Red letters indicate highly conserved columns (exact conservation of the amino acid identity), and blue indicates conservation of the amino acid properties (basic, acid, neutral) but not identities. Gray represents areas of no conservation. **(D)** Fluorescence image of CRISPR-tagged UNC-17::mKate2 (*ot907*) animal. Scale Bar = 10μm. **(E)** Schematic of UNC-17 predicted topology (magenta) along the membrane (gray). Green represents the locations of fluorescent tags tested for thrashing assays. Alleles, either extrachromosomal arrays expressed in *unc-17* (*e245*) mutants or CRISPR knock-in strains in the *unc-17* locus, are listed based on where the fluorescent tag was inserted. **(F)** *unc-17(e245)* mutant animals barely swim (1.7 ± 2). Over-expression of untagged P*unc-17*::UNC-17 (*olaEx5703*) (114.8 ± 18) and N-terminus-tagged UNC-17::GFP (*olaEx5400*) (91.1 ± 22) swim like wild-type animals (99.4 ± 20). GFP-tag between TM6-7 (*olaEx5704*) (34.3 ± 19) or CRISPR-insertion of C-terminus mKate2 tag (*ot907*) (77.7 ± 17) leads to reduced swimming behavior when compared to wild-type animals. Insertion of GFP-FLP-on cassette at the N-terminus end (s*yb7251*) (82.9 ± 16) or insertion of the GFP-FLP-on cassette at the C-terminus (*ola503*) (101.9 ± 16) thrash as well as wild-type animals (109 ± 25). We are unsure as to the phenotypic differences of *ola503* allele and *ot907* in our thrashing assays, but note that in addition to the use of different fluorescent proteins, each allele also employs distinct linker sequences between UNC-17 and the fluorescent protein (new Figure S6). *syb7882* allele, which employes mRuby3 with a similar linker to *ola503*, also does not display defects in the thrashing assay (Figure S6). Mean ± Standard Deviation. Kruskal-Wallis test with Dunn’s multiple comparison post hoc test. **** represents p<0.0001; ** represents p<0.01; * represents p<0.05; and NS means “not significant”.

**Figure S6.**
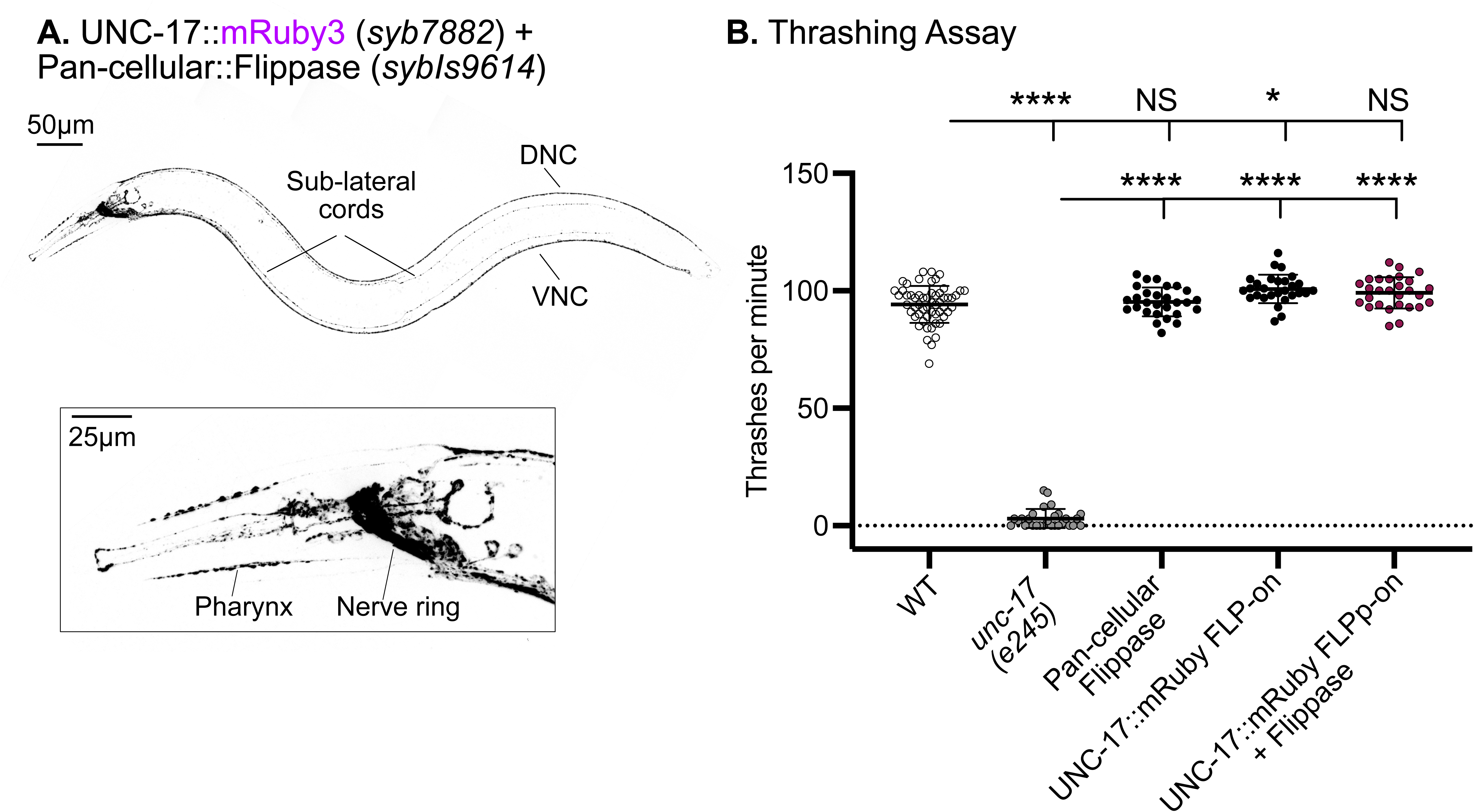
**(A)** (Top) Fluorescent image of an adult worm expressing endogenously labeled UNC-17::mRuby3 in all cells (P*eft-3*::Flippase). Scale Bar = 50μm. (Bottom) Zoom-in area of the head shows UNC-17 expressed in the nerve ring. DNC = Dorsal Nerve Cord, VNC = Ventral Nerve Cord. Scale Bar = 25μm. **(B)** *unc-17(e245)* mutant animals (3 ± 4.1) thrash significantly less than wild-type animals (94 ± 7.8). UNC-17::mRuby3 FLP-on (*syb7882*) animals that express pan-cellular::Flippase (99 ± 6.6) (*sybIs9614*) or animals that only express pan-cellular::Flippase (95 ± 6.2) are indistinguishable from wild-type animals in their thrashing behavior. UNC-17::mRuby FLP-on animals (100 ± 6.1) thrash significantly more than wild-type animals. Mean ± Standard Deviation. Kruskal-Wallis ANOVA test with Dunn’s multiple comparisons post hoc test. **** represents p<0.0001; * represents p<0.05; and NS means “not significant”.

**Figure S7.**
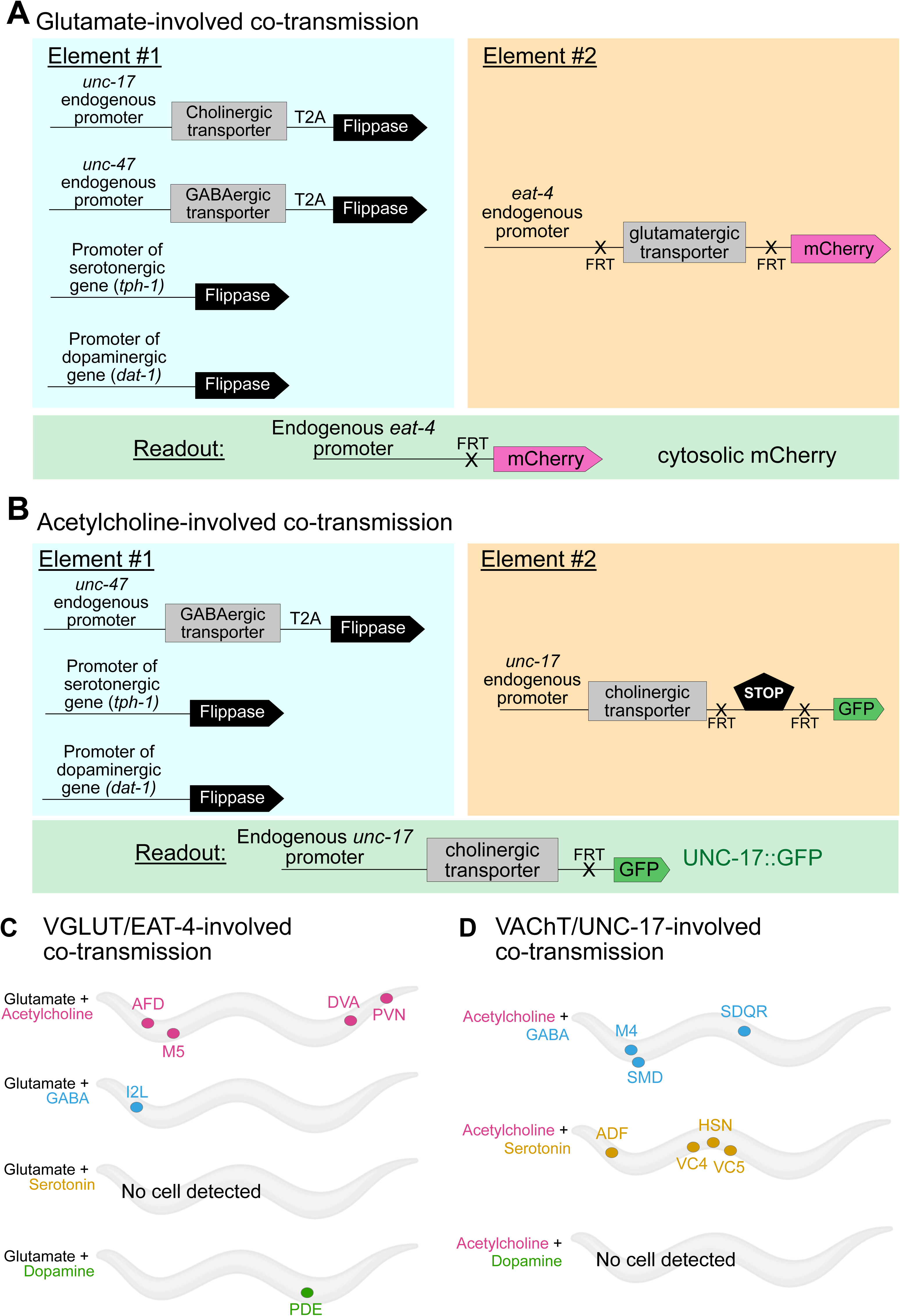
**(A)** To track glutamate-involved co-transmission, we repurposed the *eat-4* conditional KO strain (*kySi76 kySi77*) (Lopez-Cruz et al., 2019) where the *eat-4* gene coding sequence is flanked by two FRT sites and followed by cytosolic mCherry (Element #2, orange box). Crossing this line with a panel of flippase drivers (Element #1, blue box) results in activation of cytosolic mCherry only in cells were both elements were co-expressed (Readout, green box). P*dat-1*::Flippase and P*tph-1*::Flippase driver strains were made available from (Muñoz-Jiménez et al., 2017). **(B)** To track acetylcholine-involved co-transmission, we repurposed the unc-17::GFP conditional KI strain (*ola503*) (Figure 3B) for which, after the *unc-17* gene coding sequence, there are FRT sites flanking the STOP codon and 3’ UT, followed by GFP (Element #2, orange box). Crossing this line with a panel of flippase drivers (Element #1, blue box) results in the protein fusion of UNC-17 with GFP in cells where both elements were co-expressed (Readout, green box). **(C-D)** Systematic mapping shows neurons with co-transmission of **(C)** glutamate in combination with acetylcholine, GABA, and dopamine. **(D)** Co-transmission of acetylcholine is found in combination with GABA and serotonin.

**Figure S8.**
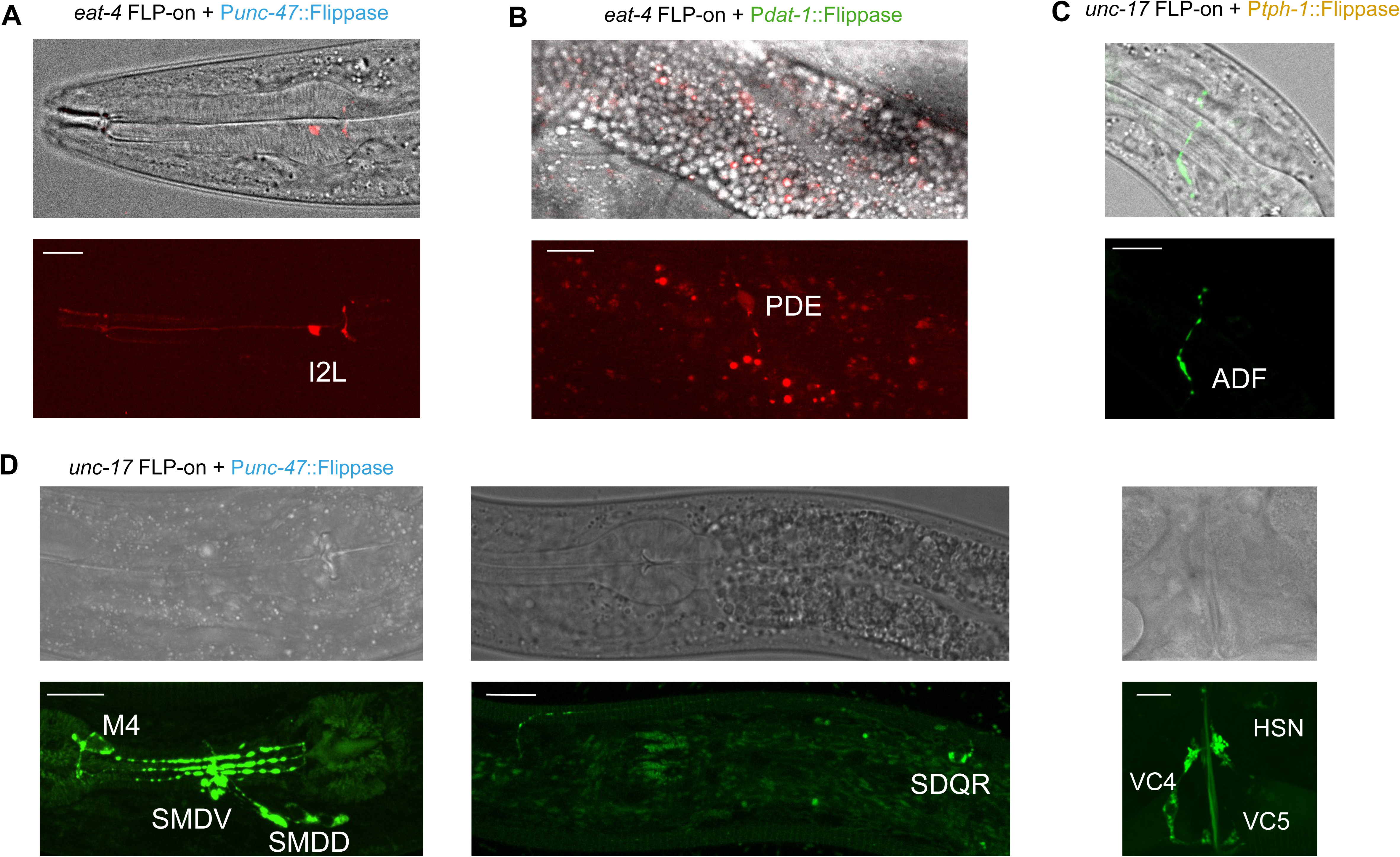
Live-imaging of co-expression strategy of the *eat-4* conditional knockout strain (*kySi76 kySi77*) with **(A)** endogenous P*unc-47*::UNC-47::T2A::Flippase and **(B)** P*dat-1*:: Flippase driver. Live-imaging of co-expression strategy of the *unc-17* conditional GFP knock-in strain (*kySi76 kySi77*) with **(C)** P*tph-1*:: Flippase driver, and with **(D)** endogenous P*unc-47*::UNC-47::T2A::Flippase. (Top) DIC images. (Bottom) Fluorescence imaging. All scale bars = 10μm.

**Figure S9.**
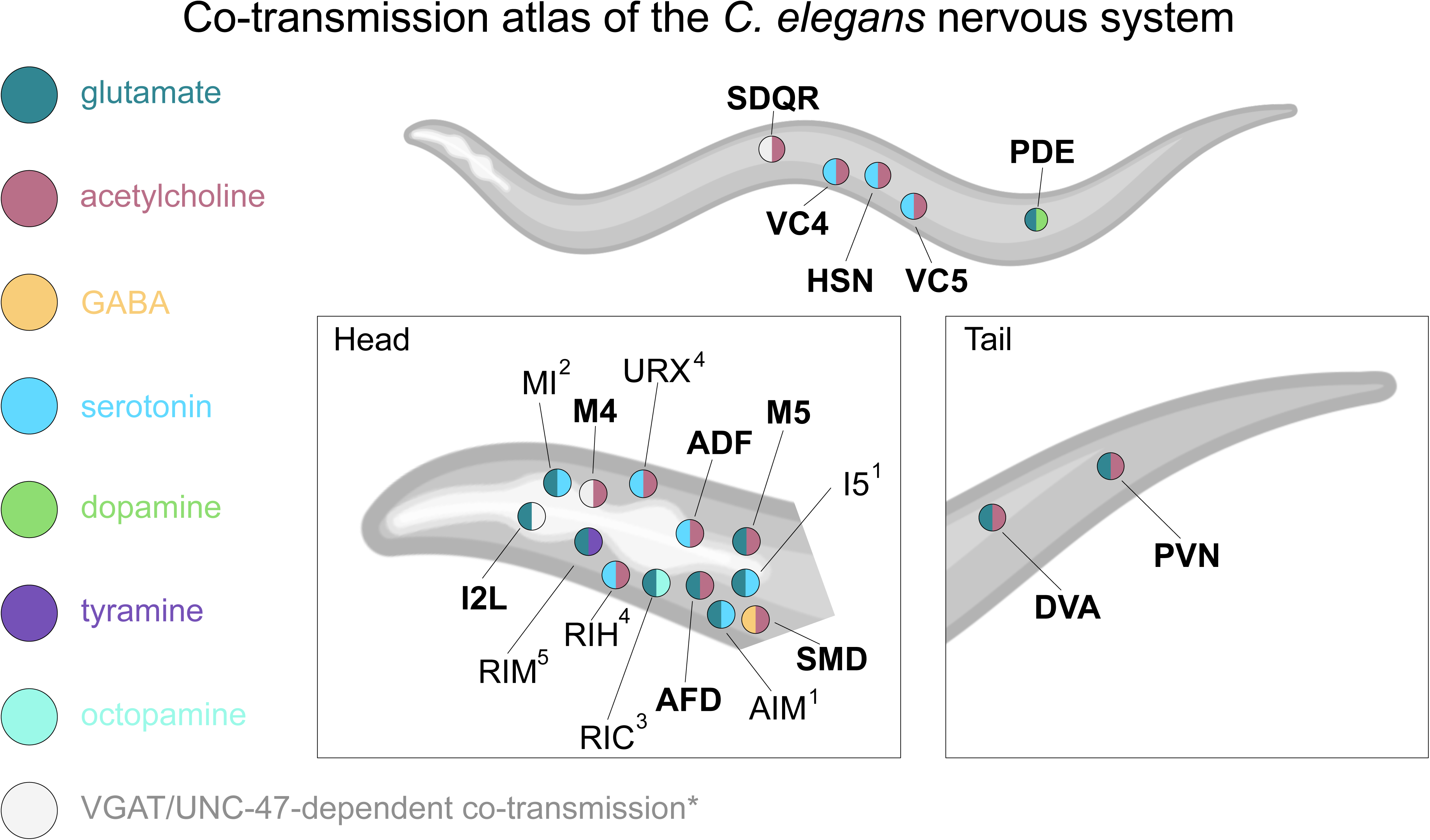
Our results (bold) are consistent and extend findings from previous studies in the field (See Table 3 and numbered references below). Here, we present an atlas of co-transmission in the *C. elegans* nervous system. Co-transmitter neurons are present in the (i) head, mid-body region and (ii) tail of the animal. *Note that I2L, M4 and SDQR express UNC-47/VGAT but do not stain positive for GABA staining (Gendrel et al., 2016) nor express genes required for GABA synthesis or uptake. ^1^ Jafari et al. (2011), Serrano-Saiz et al. (2013), and Wang et al. (2024) ^2^ Serrano-Saiz et al. (2013), and Wang et al. (2024) ^3^ Reilly et al. (2022), and Wang et al. (2024) ^4^ Jafari et al. (2011), and Wang et al. (2024) ^5^ Alkema et al. (2005), Serrano-Saiz et al. (2013), and Wang et al. (2024)

**Figure S10.**
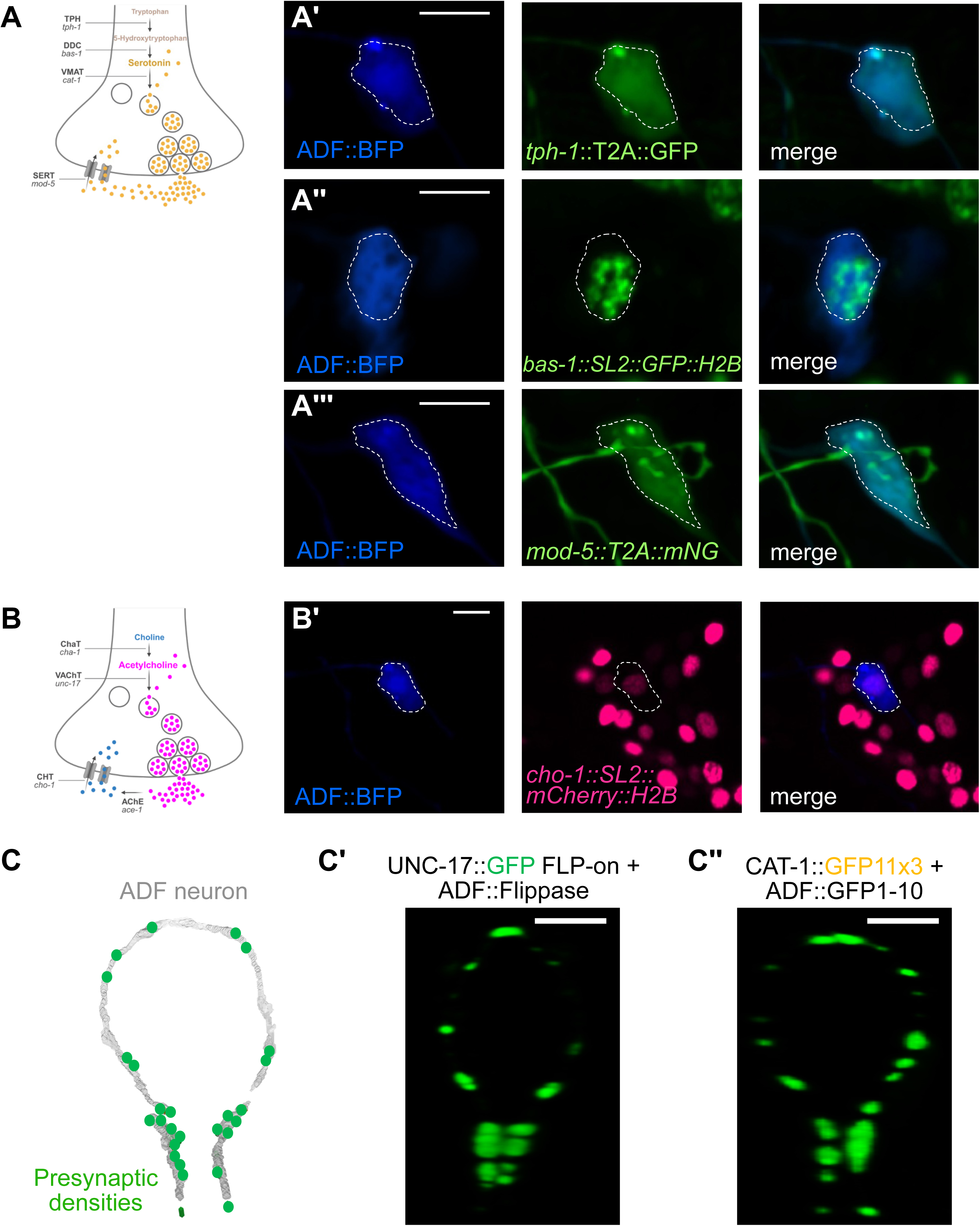
**(A)** Schematic of genes required for serotonergic identity. Mammalian (bold) and *C. elegans* (italics) homologue genes are listed. Detectable expression of **(A’)** *tph-1,* **(A’’)** *bas-1* and **(A’’’)** *mod-5* in ADF neuron (blue). Scale Bar = 5 μm. **(B)** Schematic of genes required for cholinergic identity. Mammalian (bold) and *C. elegans* (italics) homologue genes are listed. Detectable expression of **(B’)** *cho-1* in ADF neuron (blue). Scale Bar = 10 μm. **(C)** Presynaptic densities (green) on ADF axons as determined by serial electron microscopy reconstructions (White et al., 1986) (image generated with NeuroSC (Koonce et al., 2025). **(C’)** 3D rendering of live fluorescence imaging of endogenous UNC-17::GFP and **(C’’)** CAT-1::GFP specifically in ADF neurons. Scale bar = 10 μm.

